# Temporal Dynamics of Cortical State Plasticity Following Adult Vision Loss

**DOI:** 10.64898/2026.04.21.719808

**Authors:** Ismaël Djerourou, Maurice Ptito, Matthieu P. Vanni

**Affiliations:** École d’optométrie, Université de Montréal, Montréal, Québec H3C 3J7, Canada; Centre interdisciplinaire de recherche sur le cerveau et l’apprentissage (CIRCA), Université de Montréal, Montréal, Québec H3C 3J7, Canada

## Abstract

The adult brain retains a capacity for adaptation, yet the long-term sequence of plasticity remains poorly understood. To address this, we performed longitudinal mesoscopic calcium imaging of the dorsal cortex in spontaneously behaving adult mice before and after bilateral enucleation (BE). We uncovered behaviour-dependent and spatially distinct changes in cortical activity organised into two sequential, overlapping windows. An early window, spanning from day 1 to week 3–5, was marked by reduced activity in visual and retrosplenial cortices during locomotion, with movements suppressing rather than enhancing activity, as is typically observed. A second, delayed window (week 1 to week 7–10, peaking around 3 weeks) was characterised by increased slow-wave activity in primary visual cortex (V1) and lateromedial visual area (LM), indicative of heightened excitability. Consistently, activity increased—rather than decreased—at the transition from locomotion to quiescence. In parallel, we observed a rapid, long-lasting, and behaviour-dependent reorganisation of cortical network structure. Together, these findings demonstrate that cortical states in the adult brain are dynamic and can undergo substantial, months-long plasticity. The temporal and spatial dissociations between windows identified suggest distinct plasticity mechanisms in the circuit regulating cortical states. Therefore, our study provides a spatiotemporal framework to guide future investigations into the mechanisms of plasticity in the mature brain.

## Introduction

The mature brain retains a remarkable capacity for adaptation beyond the closure of critical periods of plasticity (Gilbert & Li, 2012; Hübener & Bonhoeffer, 2014; H.-K. Lee & Whitt, 2015). Years after vision loss in adult humans, the visual cortex is recruited by many non-visual tasks, reflecting robust cross-modal reorganisation of deprived visual areas (Kupers & Ptito, 2014; Paré et al., 2023). However, the sequence of plasticity mechanisms underlying this long-term adaptation remains poorly understood.

In adult mice, the early stages of this process in the primary visual cortex (V1) have been partially characterised. Within 1–2 days following visual deprivation, excitatory activity is restored through homeostatic plasticity mechanisms (W. Wen & Turrigiano, 2024), involving a disengagement of inhibitory neurons and a strengthening of excitatory synapses onto pyramidal neurons, thereby re-establishing the excitation–inhibition balance (Barnes et al., 2015; J. L. Chen et al., 2011; Keck et al., 2011, 2013). At a much later stage—approximately seven weeks after deprivation—post-mortem analyses reveal the emergence of cross-modal plasticity driven by whisker inputs (Van Brussel et al., 2011). Together, these findings point to a substantial temporal gap between rapid homeostatic adjustments and the onset of cross-modal reorganisation, spanning several weeks during which the dynamics of cortical adaptation remain largely unknown.

To address this gap, we performed longitudinal mesoscopic calcium imaging (Cardin et al., 2020; Couto et al., 2021; Silasi et al., 2016; Vanni et al., 2017) in head-fixed behaving mice before and after bilateral enucleation (BE). During initial data exploration, we observed an unexpected shift in V1 activity across behavioural states following BE. In sighted mice at baseline, V1 activity was lower during quiescence than during locomotion, consistent with well-established state-dependent modulation of cortical activity (Polack et al., 2013; Poulet & Crochet, 2019; Reimer et al., 2014; Stringer et al., 2019; Vinck et al., 2015; West et al., 2022). Strikingly, in the same animals, three weeks after BE, this relationship reversed, with V1 activity becoming higher during quiescence than during locomotion. This unexpected reversal motivated a systematic investigation of behavioural state as a key dimension of cortical plasticity—an aspect that has been largely overlooked in longitudinal studies of adult plasticity.

We identified months-long, behaviour-dependent plasticity of cortical activity following adult vision loss. This process unfolded in two distinct temporal windows. First, during an early phase spanning from day 1 to approximately 3–5 weeks after bilateral enucleation (BE), locomotion exerted an inverted effect on cortical activity, suppressing rather than activating the visual and retrosplenial cortices. Second, during a later phase extending from week 1 to approximately 7–10 weeks, the primary visual cortex (V1) and the lateromedial visual area (LM) exhibited increased slow-wave activity during quiescence. In parallel with these state-dependent changes, we observed a rapid and persistent reorganisation of cortical networks that remained tightly linked to behavioural state. Together, these findings demonstrate that the adult brain undergoes prolonged plasticity of cortical states following vision loss.

## Results

### Longitudinal mesoscopic calcium imaging during spontaneous behaviour

We recorded dorsal cortical calcium activity in adult Thy1-jRGECO1a mice (Dana et al., 2018) during head-fixed spontaneous behaviour up to 10 weeks following BE. Mice were implanted with a chronic imaging chamber, enabling optical access to most dorsal cortical regions (Silasi et al., 2016) (Fig. 1a; see Methods). Mesoscopic calcium imaging was performed by illuminating the cortex with a lime LED to excite jRGECO1a and recording red fluorescence at 30 Hz with a sCMOS camera (Fig. 1a). Images were segmented using the Allen Mouse Brain Atlas (Oh et al., 2014) aligned to functional visual maps (and somatosensory maps when available), and pixel values were averaged within an area around the centroid of each region. Calcium signals were corrected for hemodynamic absorption artefacts using red and green reflectance imaging (See Methods) (S. Shahsavarani et al., 2023; Wang et al., 2024). Behaviour was monitored using infrared video recordings at 30 Hz, and movement (motion energy) was extracted from a manually defined region of interest using Facemap (Syeda et al., 2024) (Fig. 1a; see Methods).

**Figure 1.**
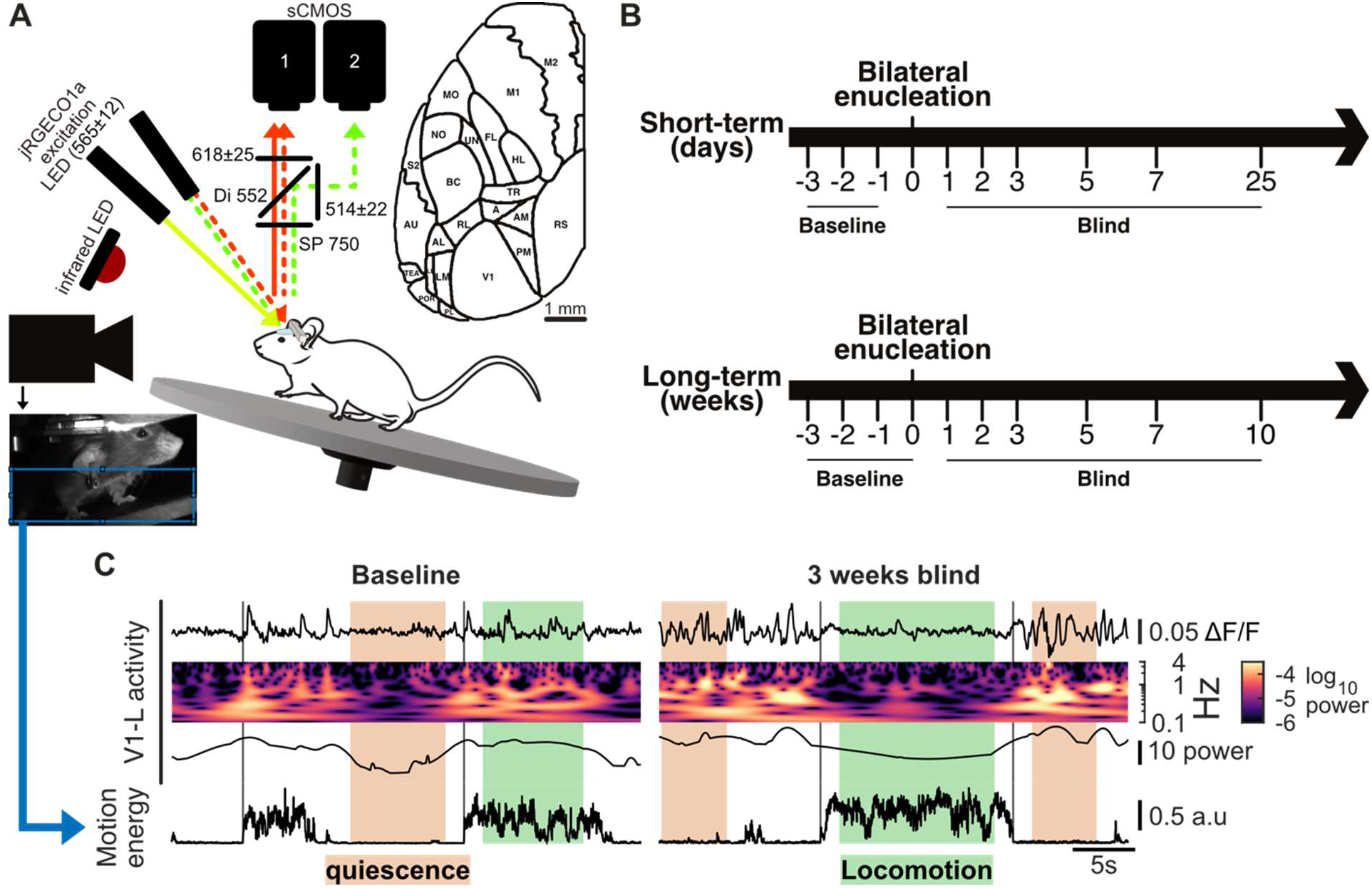
Setup and longitudinal protocols. **a**, Setup for mesoscale calcium imaging of the mouse dorsal cortex during head-fixed spontaneous behaviour. Straight lines indicate the excitation and emission light for calcium imaging. Dashed lines indicate the light used for hemodynamic correction of the calcium signal. An example view from the behavioural camera with the ROI used to compute the motion energy is shown. Mouse image from Scidraw.io (doi.org/10.5281/zenodo.3926135). **b**, Longitudinal imaging protocols. The top panel shows the short-term protocol, with sessions spanning days after BE. The bottom panel shows the long-term protocol, with sessions spanning weeks. **c**, Example data from two sessions in the same mouse from the long-term protocol. The first row shows the hemodynamic-corrected calcium signal (ΔF/F) from the left V1. The second row shows the continuous wavelet transform (CWT) of the signal (cropped to the window illustrated in the first row) within the 0.1–4 Hz band. The third row displays the maximum power over time extracted from the CWT. The last row shows the motion energy, normalised between 0 and 1. Behavioural segmentation into quiescence (orange), locomotion (green), and state transitions (vertical lines), used in the analysis, is overlaid across all rows.

This approach was applied in both short-term (1–25 days post-BE) and long-term (1–10 weeks post-BE) longitudinal protocols (Fig. 1b).

### Inversion in the relationship between movement and visual cortical activity

During data exploration, we observed that V1 activity in a session recorded three weeks after BE was higher during quiescence than during locomotion—the opposite of both the expected pattern and that observed in the same mouse at baseline (Fig. 1c). This unexpected finding prompted us to systematically examine how the relationship between movement and cortical activity changes following BE. Using the same sessions, we computed the Spearman correlation between V1 activity and motion energy. At baseline, the correlation was positive (p < 0.05), indicating that increased movement was associated with higher activity. In contrast, three weeks after BE, the correlation became negative (p < 0.05), indicating that increased movement was associated with reduced activity (Fig. 2a). Extending this analysis across mice, cortical regions, and time points, we identified a significant effect of BE on the relationship between movement and activity in V1 as well as in higher visual areas, including lateromedial (LM), anterolateral (AL), and posteromedial (PM) areas, across both experimental protocols (Fig. 2b). In V1, correlations decreased rapidly from day 1 post-BE, shifting from positive at baseline to negative values. From week 3 onward, correlations gradually increased, but remained below baseline levels even at week 10 (Fig. 2c, Table S1). Together, these results demonstrate that visual deprivation induces a sustained reorganisation of movement-related cortical states, leading to an inversion of the canonical relationship between locomotion and visual cortical activity.

**Figure 2.**
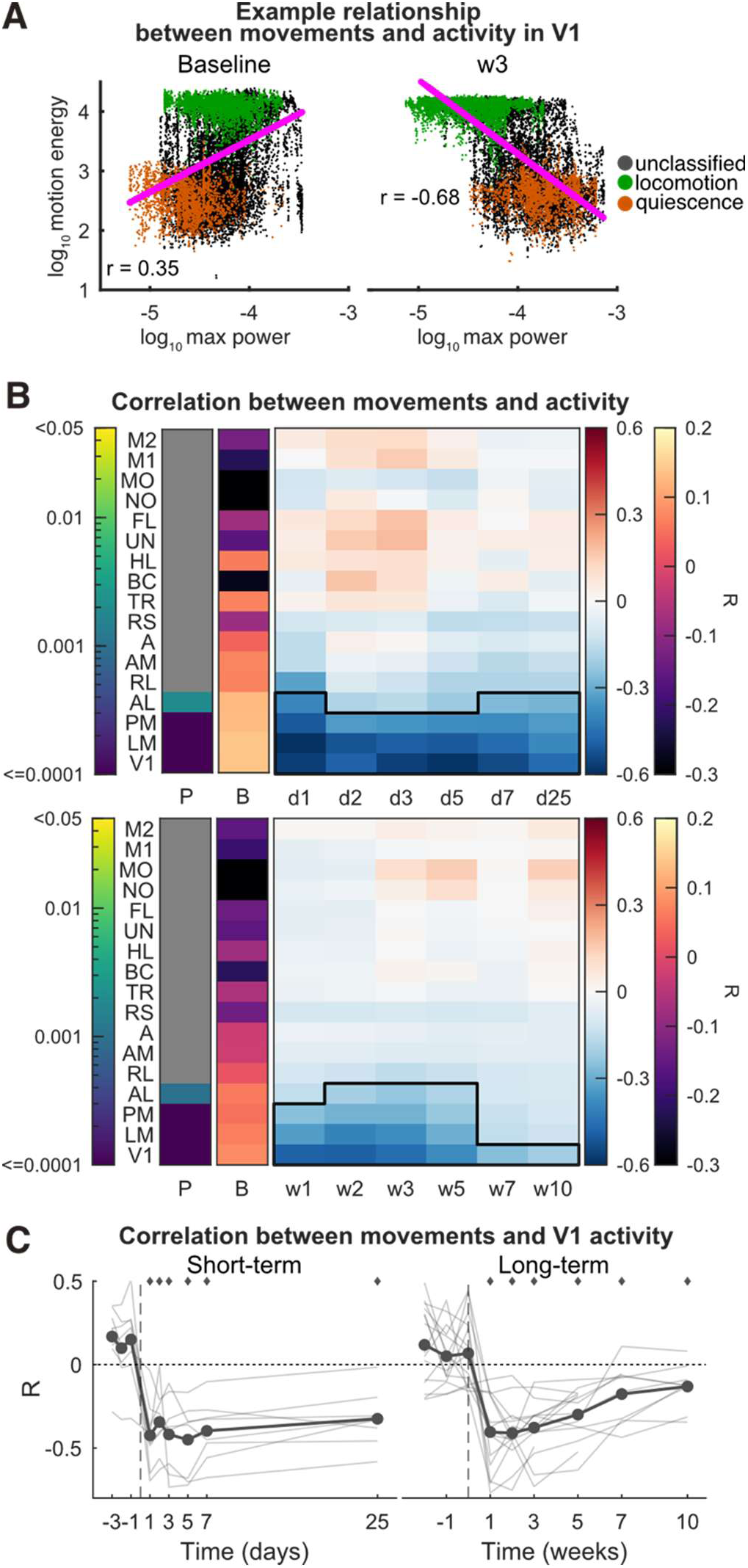
Inversion in the relationship between movements and visual cortex activity. Scatter plots of the motion energy versus maximum power in left V1 (V1-L) from two sessions of the same mouse: baseline and 3 weeks after BE. Each dot is a data point from the timeseries of the example session. They are colour-coded by behavioural state. Data points that are not associated with a behavioural state have been labelled as unclassified. Magenta lines show linear regression fits, with corresponding Spearman correlation coefficients. **b** Spearman correlations between the motion energy and maximum power across cortical regions and time. *P* matrices (P) indicate regions with significant effects of time in a linear mixed-effects ANOVA, FDR-adjusted across regions (*p* < 0.05). The colour bar indicates log10(*p*). Baseline matrices (B) show the correlation coefficients averaged across mice and sessions. Post-BE matrices show the difference from baseline. The black outline indicates time points that are significantly different from baseline (*p* < 0.05, post hoc FDR-adjusted). **c** Spearman correlation between the motion energy and maximum power in V1. Thin lines represent individual mouse data. Diamonds mark significant time points relative to baseline (*p* < 0.05, post hoc FDR-adjusted). *(**b,c)** See Table S1 for details about the statistics. See Figure S1 for the number of mice per time point*.

### Sequential and overlapping windows of behaviour-defined plasticity

To characterise the temporal dynamics of behaviour-related cortical activity following bilateral enucleation (BE), we segmented mouse behaviour into quiescence and locomotion epochs based on motion energy (Fig. 1c). Transitions were identified using a change-point detection algorithm (Lohani et al., 2022; Vinck et al., 2015), and included epochs ≥10 s and separated by 3 s from transitions. Overlaying behavioural states onto activity distributions revealed a clear segregation of V1 activity between locomotion and quiescence (Fig. 2a). For each epoch, we computed the power spectral density and quantified activity as the maximum power within the 0.2–4 Hz band.

During quiescence, BE significantly altered activity in V1 and the lateromedial area (LM) across both protocols (Fig. 3a). V1 activity exhibited a transient increase at day 1, recovered by day 2, and then progressively increased again from week 1 onward, remaining elevated relative to baseline at week 10. Spectral analysis revealed that this increase was driven by enhanced slow-wave activity: power in the 0.2–4 Hz range increased at day 1 (peaking near ∼1 Hz), returned to baseline at day 2, and rose again from week 1 to 10, with peak changes initially in the 0.4–0.6 Hz range and shifting toward lower frequencies over time (Fig. 3b, Fig. S2a). These results indicate a sustained, slow-wave–dominated state during quiescence following BE.

**Figure 3.**
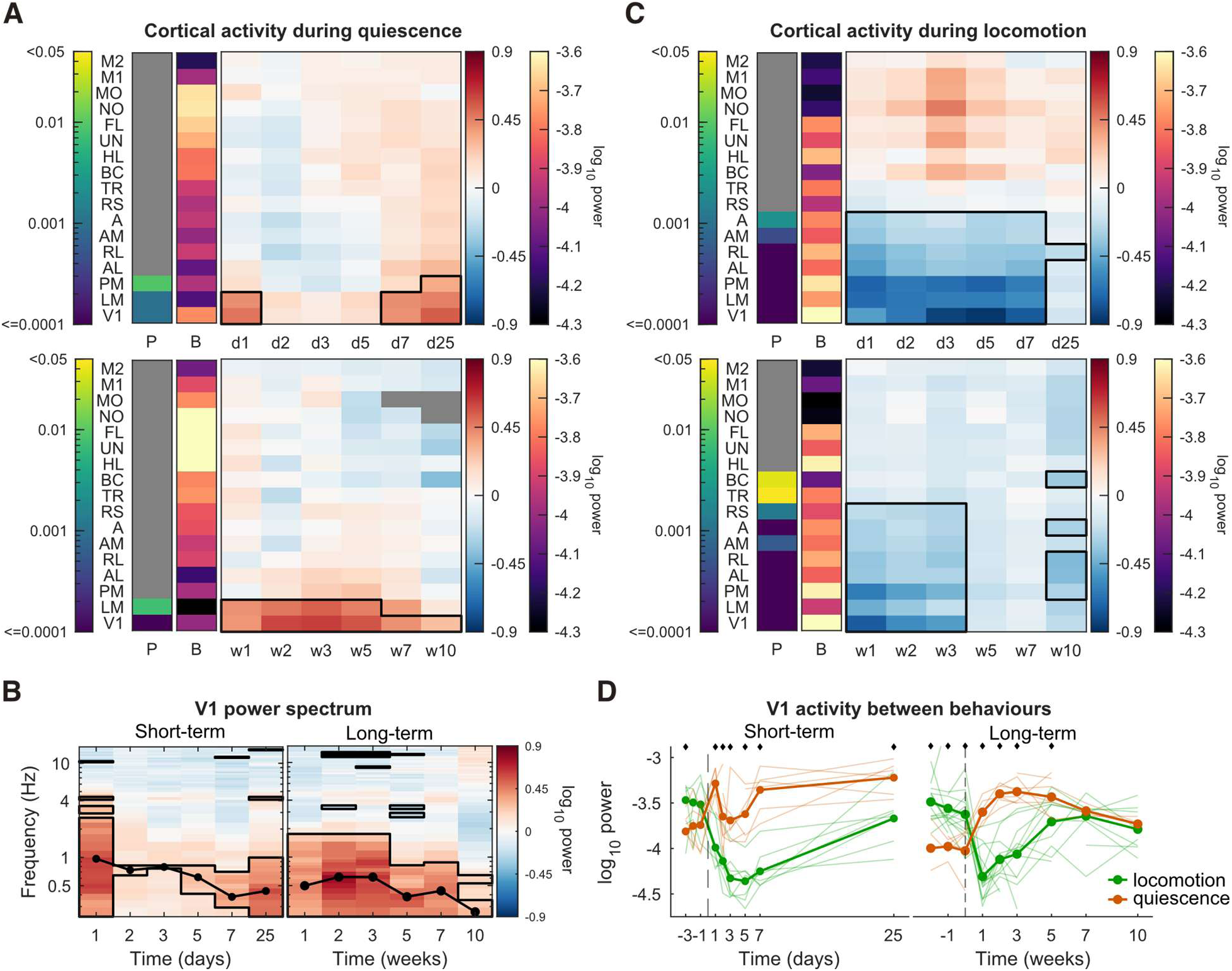
Sequential and overlapping windows of behaviour-defined plasticity. **a** Maximum power during quiescence across cortical regions and time. P matrices (P) indicate regions with significant effects of time in a linear mixed-effects ANOVA, FDR-adjusted across regions (p < 0.05). The colour bar indicates log10(p). Baseline matrices (B) show the maximum power averaged across mice and sessions. Post-BE matrices show the difference from baseline. The black outline indicates time points and regions that are significantly different from baseline (p < 0.05, post hoc FDR-adjusted). **b,** Frequency power spectrum of V1 activity during quiescence, shown as the difference with baseline. The black outline indicates time points and frequencies that are significantly different from baseline (p < 0.05, post hoc FDR-adjusted). **c**, Same as **a**, but for locomotion. **d**, Maximum power over time during quiescence and locomotion. Thin lines represent individual mouse data. Diamonds mark significant time points with differences between states (*p* < 0.05, post hoc FDR-adjusted). *(**a–d**) See Table S1 for details about the statistics. See Figure S1 for the number of mice at each time point*.

In contrast, during locomotion, BE led to a marked suppression of activity in visual and retrosplenial cortices (Fig. 3c). In V1, activity was significantly reduced from day 1 to week 3, reaching a minimum around day 5 before recovering to baseline by week 5. Power spectra showed a broad reduction across frequencies, consistent with a global decrease in activity (Fig. S2c,d).

Direct comparison between behavioural states revealed a reversal of the canonical relationship between locomotion and quiescence. At baseline, V1 activity was higher during locomotion, whereas after BE, locomotion-associated activity became significantly lower than quiescence until approximately week 7, after which no significant difference remained (Fig. 3d). This pattern is consistent with the correlation analyses (Fig. 2) and indicates a state-dependent inversion of cortical activity dynamics.

Importantly, these effects were not explained by changes in behavioural structure. Although variability in state duration and epoch number was observed across animals (Fig. S3a,b), these measures were stable on average and did not account for the observed neural changes. Motion energy levels during locomotion and quiescence also remained largely unchanged relative to baseline (Fig. S3c,d), aside from a transient increase at day 7.

Together, these findings reveal two sequential and partially overlapping windows of plasticity in the visual cortex following vision loss: an early phase (day 1 to week 3) characterised by locomotion-related suppression of activity, and a later phase (week 1 to week 10) marked by enhanced slow-wave activity during quiescence. These results demonstrate prolonged, state-dependent reorganisation of cortical dynamics after sensory deprivation.

### Movement initiation suppresses posterior cortical activity after BE

As stated above, the analysis of activity during quiescence and locomotion revealed two sequential, overlapping windows of plasticity. We next asked whether these changes in activity were present at transitions between behavioural states. Transition times were identified using a change-point detection algorithm applied to motion energy, and for each transition, we computed ΔF/F₀ relative to the pre-transition baseline.

At baseline, the transition from quiescence to locomotion (locomotion onset) was marked by a rapid increase in motion energy accompanied by a robust increase in V1 activity, which remained elevated for several seconds (Fig. 4a). A similar response was observed in other cortical regions, including the hindlimb (HL) area of the primary somatosensory cortex (Fig. S4a). In contrast, from day 1 after bilateral enucleation (BE), locomotion onset failed to elicit this activation in V1, while responses in HL remained largely unchanged (Fig. 4a, Fig. S4a).

**Figure 4.**
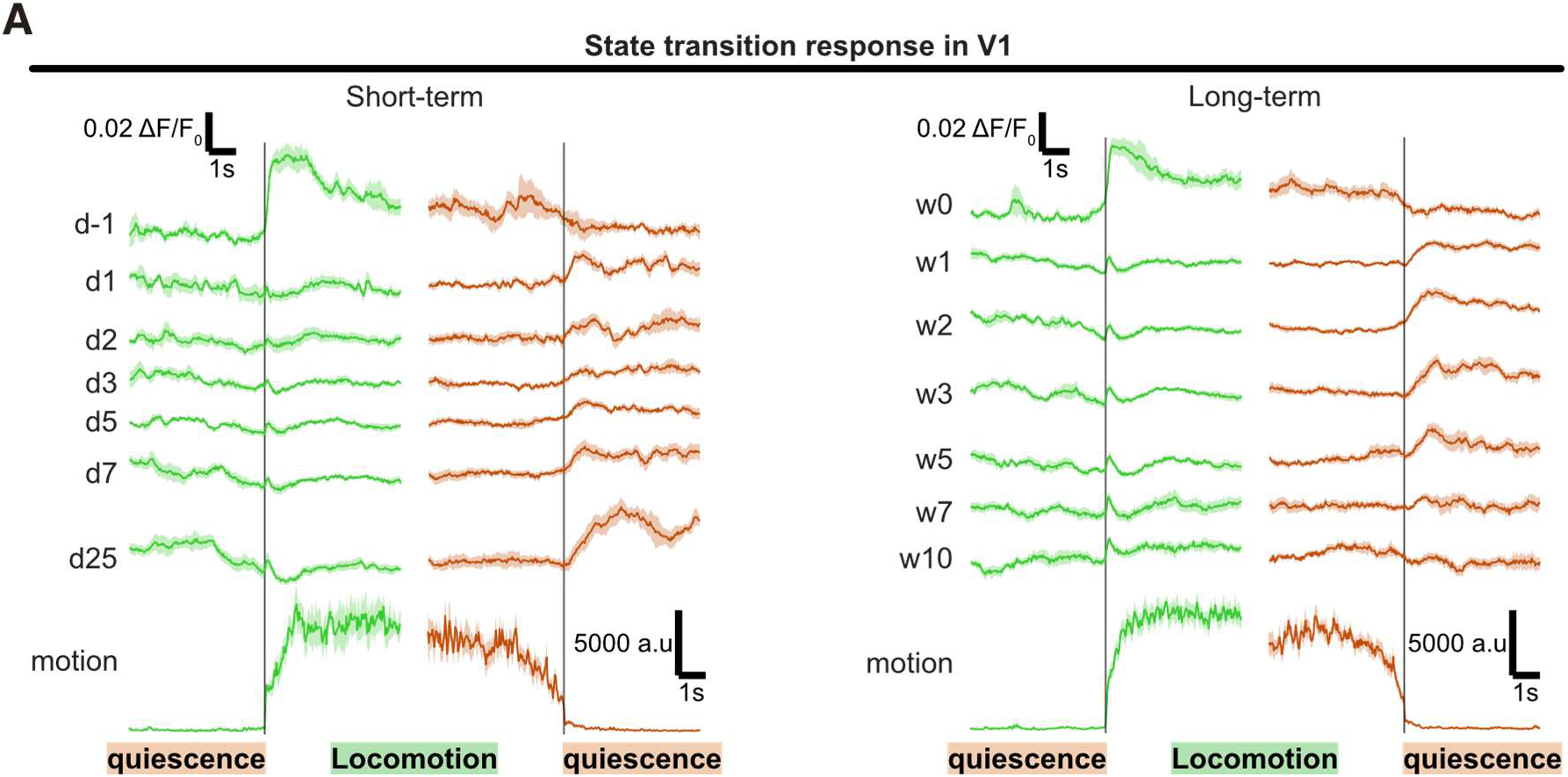
V1 activity at the onset and offset of locomotion. a, Mouse averaged ΔF/F_0_ response in V1 time locked on the locomotion onset and offset at one baseline and every time point after BE. The last row shows the motion energy time-locked on the transitions from the baseline session. The shaded areas represent the SEM.

To quantify these effects longitudinally, we measured the mean ΔF/F₀ within a 1 s window following locomotion onset. Across both protocols, activity in visual and retrosplenial cortices was consistently reduced after BE, with most visual areas—including V1—showing suppression from day 1 through week 10 (Fig. 5a). This reduction persisted longer than the suppression observed during sustained locomotion (Fig. 3c), with only partial recovery toward baseline after week 2 and significant deficits still present at week 10 (Table S1). The suppression extended beyond the 1 s window, peaking between 0.5 and 1 s after transition (Fig. 5b), indicating that locomotion-related suppression starts immediately at movement onset and persists over an extended time course.

**Figure 5.**
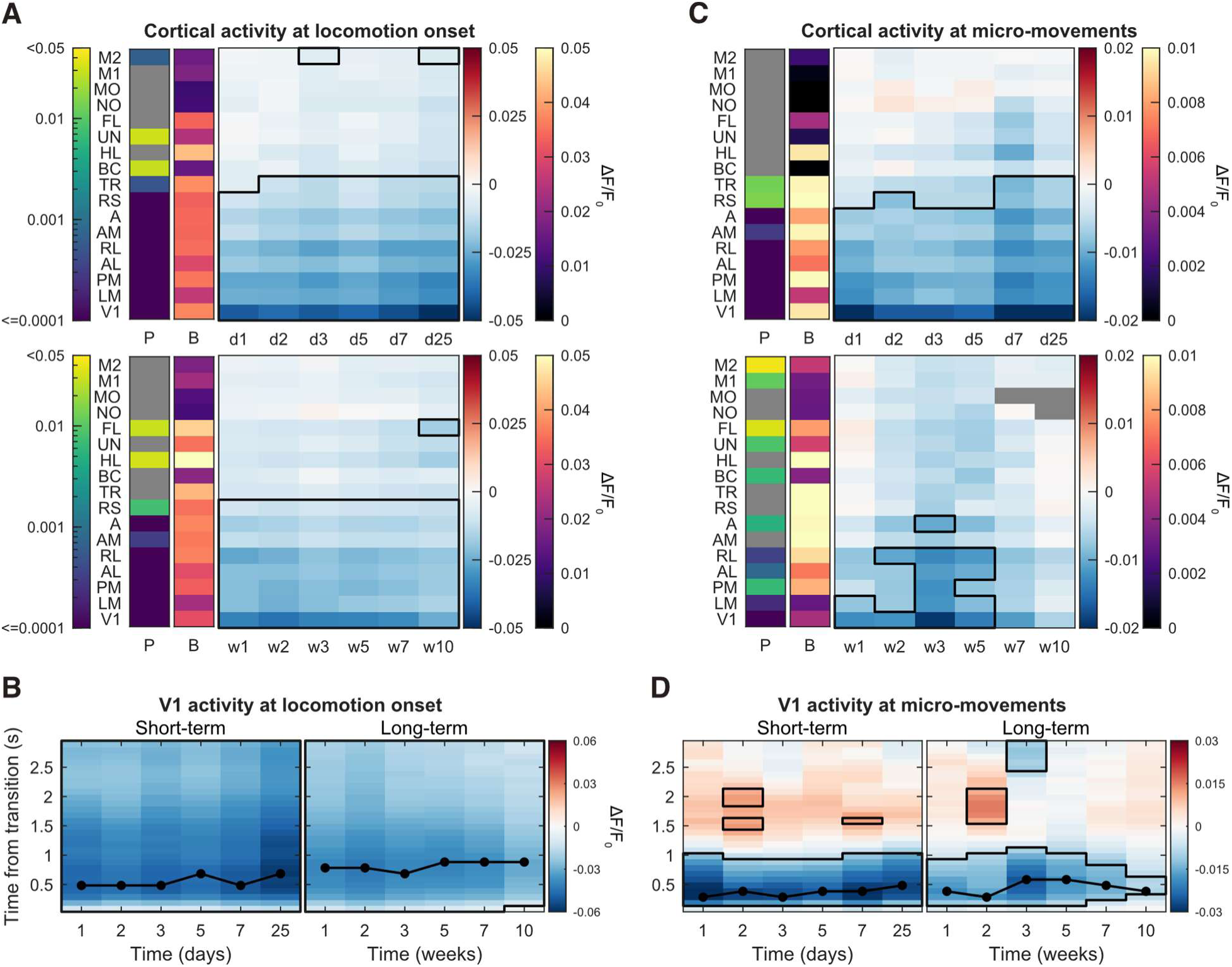
Movement initiation suppresses posterior cortical activity after BE. a, Average ΔF/F_0_ within a 1s window following locomotion onset across cortical regions and time. P matrices (P) indicate regions with significant effects of time in a linear mixed-effects ANOVA, FDR-adjusted across regions (p < 0.05). The colour bar indicates log10(p). Baseline matrices (B) show the ΔF/F_0_ averaged across mice and sessions. Post-BE matrices show the difference from baseline. The black outline indicates time points and regions that are significantly different from baseline (p < 0.05, post hoc FDR-adjusted). **b** V1 activity following locomotion onset is shown as the difference with baseline. The black curve indicates the time from transition with maximum reduction in activity. The black outline indicates protocol and transition time points that are significantly different from baseline (p < 0.05, post hoc FDR-adjusted). **c** same as **a** but for micro-movements. **d** same as **b** but for micro-movements. *(**a–d**) See Table S1 for details about the statistics. See Figure S6 for the number of mice at each time point*.

We next asked whether this suppression was specific to locomotion or generalised to other forms of movement. To address this, we analysed brief, low-amplitude movements during quiescence (“micro-movements”; Fig. S5a). At baseline, these events elicited transient increases in activity across multiple cortical regions, including V1 (Fig. 5c, Fig. S5a). Following BE, however, visual cortical responses were selectively reduced, with V1 activity significantly suppressed from day 1 to week 5 (Fig. 5c). This suppression was temporally confined to the duration of the movement (∼1 s), peaking between 0.3 and 0.6 s after onset (Fig. 5d), and could not be explained by changes in movement amplitude, as motion energy remained stable across sessions (Fig. S5).

Together, these findings demonstrate that movements broadly shift from activating to suppressing visual cortical activity following vision loss. Notably, the temporal profile of this suppression depends on movement type, with locomotion producing stronger and more prolonged effects than micro-movements. These differences likely reflect the larger baseline activation evoked by locomotion. After BE, however, both movement types converge toward similarly reduced activity levels (Fig. S5b), indicating a generalised, state-dependent reorganisation of movement-related cortical responses over a week-long period.

### Rapid increase in visual cortical activity following the cessation of locomotion after BE

Our previous analysis of cortical activity during continuous epochs of quiescence revealed a delayed, weeks-long window of increased visual cortex excitability, characterised by higher slow-wave power (Fig. 3). Here, we ask whether this increase occurred at the transition from locomotion to quiescence. Like movement activity, we used the average ΔF/F_0_ within a 1s window following the transition time to quantify temporal changes in the longitudinal protocols. We found that activity within the visual cortex significantly increased following locomotion onset. In V1, the activity increased from day 1 to week 7, with a recovery to baseline at week 10 (Fig. 6a). There was variability between protocols about the time from the transition with maximum increase relative to baseline, being between 1.3s and 2.8s for the short-term, and between 0.9s and 1.2s for the long term (Fig. 6b). The maximum change in activity mostly occurred after the 1s window investigated before (Fig. 6a). When the maximum V1 activity within the first 3s after transition was taken instead, it revealed a trajectory of change in activity over time like the one found during quiescence. V1 activity increased at day 1, partially recovered by day 2, and increased again from day 5 to week 7 (Fig. 6c). Therefore, all measures of the activity after locomotion offset (Fig. 6), and the shape of the curves (Fig. 4), suggest a recovery to baseline at week 10.

**Figure 6.**
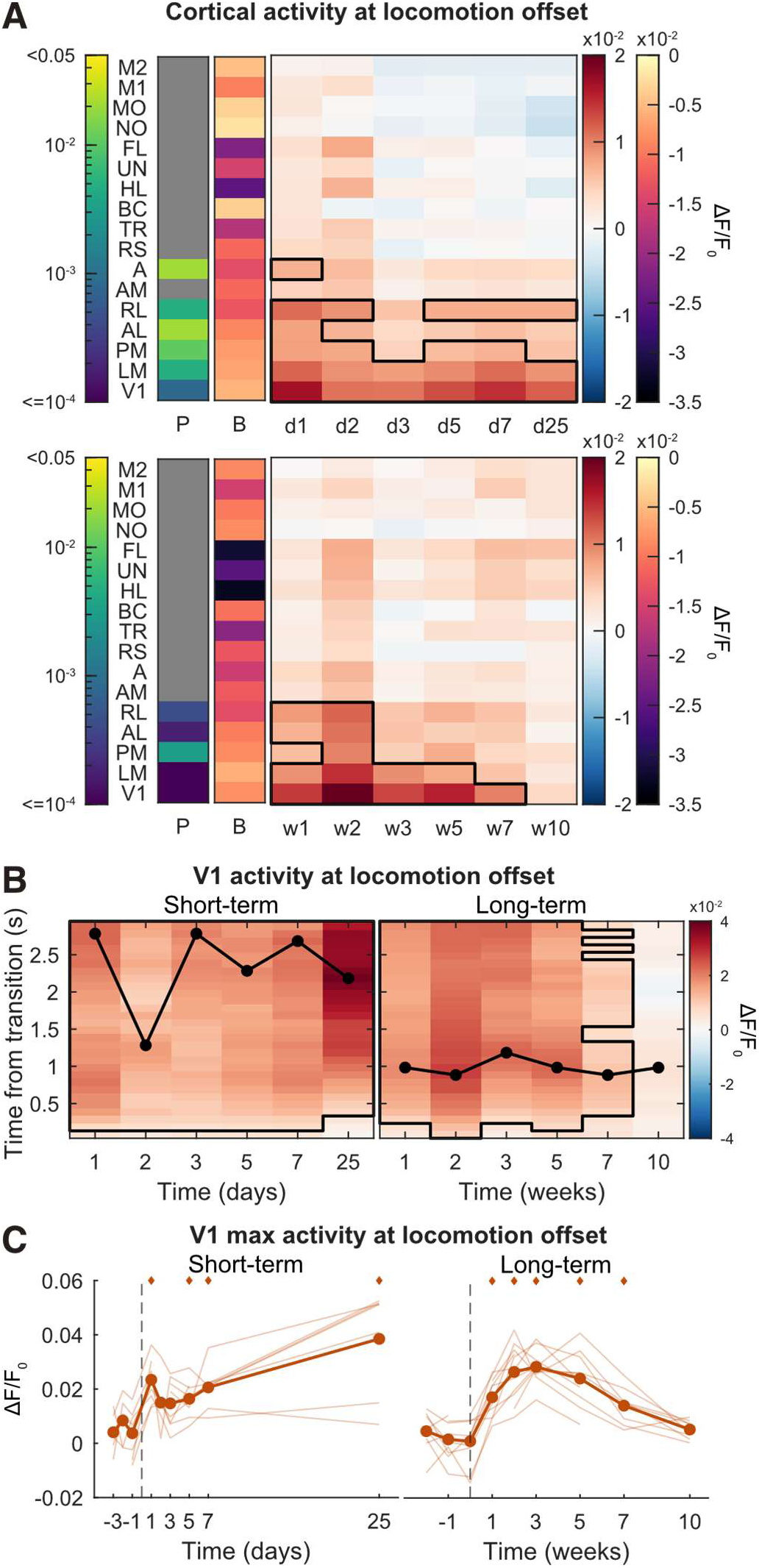
Rapid increase in visual cortical activity following the cessation of locomotion after BE. **a** Average ΔF/F_0_ within a 1s window following locomotion offset across cortical regions and time. P matrices (P) indicate regions with significant effects of time in a linear mixed-effects ANOVA, FDR-adjusted across regions (p < 0.05). The colour bar indicates log10(p). Baseline matrices (B) show the ΔF/F_0_ averaged across mice and sessions. Post-BE matrices show the difference from baseline. The black outline indicates time points and regions that are significantly different from baseline (p < 0.05, post hoc FDR-adjusted). Rapid loss of visual cortex functional connectivity after BE. **b** V1 activity following locomotion offset is shown as the difference with baseline. The black curve indicates the time from transition with maximum increase in activity. The black outline indicates protocol and transition time points that are significantly different from baseline (p < 0.05, post hoc FDR-adjusted). **c** Maximum V1 activity within a 3s window following locomotion offset. Thin lines represent individual mouse data. Diamonds mark indicates a significant difference from baseline (*p* < 0.05, post-hoc FDR-adjusted). *(**a–c**) See Table S1 for details about the statistics. See Figure S6 for the number of mice at each time point*.

We thus showed that the activity of the visual cortex rapidly increased following the transition from locomotion to quiescence, suggesting a state of increased excitability, with a delayed, weeks-long window that matched that observed during quiescence.

### Rapid, long-lasting and behaviour-dependent reorganisation of the cortical network after BE

Our regional analyses revealed two sequential, overlapping windows of behaviour-dependent plasticity with spatially distinct effects across the cortex. Such localised changes are expected to impact large-scale functional network organisation. To test this, we computed the pairwise correlation matrices from activity during spontaneous behaviour. Following BE, we observed a marked reduction in functional connectivity within the visual cortex, and between visual and retrosplenial, somatosensory, and motor regions (Fig. S7a,b). To quantify the temporal evolution of this reorganisation, we measured network similarity by correlating each time-point matrix with the first baseline matrix. This analysis revealed a sharp drop in similarity as early as day 1, which persisted throughout the 10-week recording period (Fig. S7c).

Given the behaviour-dependent changes in regional activity after BE, we next asked whether changes in cortical network organisation were similarly influenced by behavioural state. During quiescence, functional connectivity decreased within the visual cortex and between visual, retrosplenial, somatosensory, and motor regions (Fig. 7a). Network similarity analyses revealed modest and protocol-dependent effects: a transient reduction at day 2 in the short-term dataset, but a sustained decrease across all time points in the long-term dataset (Fig. 7b), indicating relatively limited reorganization in this state. Homotopic connectivity changes were restricted to V1, with a transient increase at week 2 followed by a decrease at week 10 (Fig. 7c), suggesting interhemispheric synchronisation of slow-wave activity during quiescence. Regionally, V1 intra-hemispheric correlations decreased, particularly with anterior areas, while showing a transient increase with LM between weeks 2 and 5—coinciding with the previously identified window of heightened excitability (Fig. 7d). On average, V1 correlations declined from day 1 to week 10 (Fig. 7e).

**Fig. 7.**
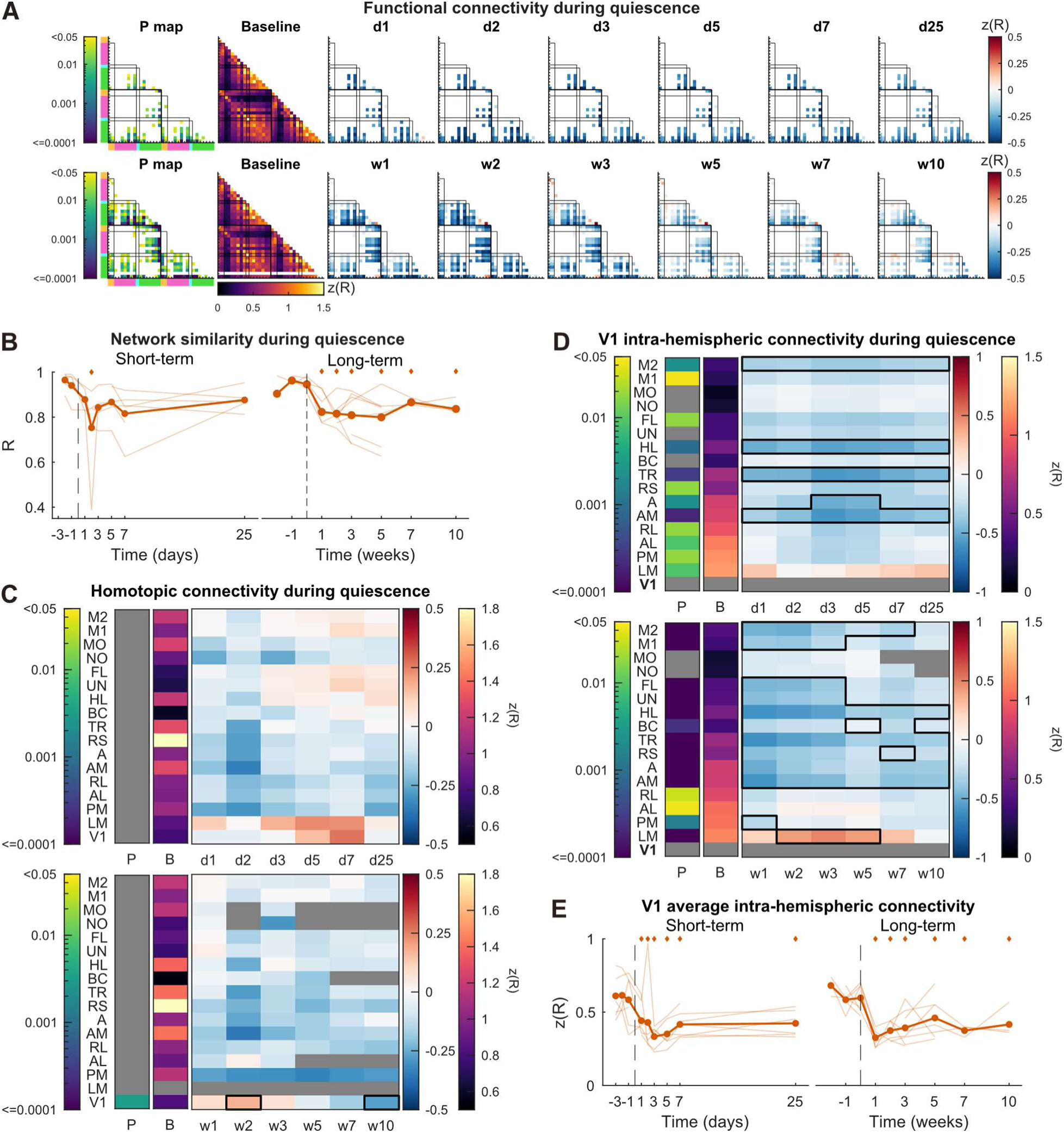
Reorganisation of the quiescent cortical network. a, Functional connectivity matrices calculated on the calcium signal (ΔF/F_0_) during quiescence using the Pearson correlation coefficient corrected with the Fisher z-transformation (z(R)). In the P matrices, only p values < 0.05 (linear mixed-effects model ANOVA, FDR-adjusted across pairs) are shown with a log10-scaled colour bar. The baseline matrices are the average correlation between mice and baseline sessions. The matrices from time points after BE show the difference from baseline, with only connections with a significant time effect shown. **b,** Network similarity. Thin lines represent individual mouse data. Diamonds mark significant time points relative to baseline (*p* < 0.05, post hoc FDR-adjusted). **c**, Homotopic correlations across cortical regions and time. *P* matrices (P) indicate regions with significant effects of time in a linear mixed-effects ANOVA, FDR-adjusted across regions (*p* < 0.05). The colour bar indicates log10(*p*). Baseline matrices (B) show the correlations averaged across mice and sessions. Post-BE matrices show the difference from baseline. The black outline indicates time points and regions that are significantly different from baseline (*p* < 0.05, post hoc FDR-adjusted). **d**, Same as **b** but for the V1 intra-hemispheric correlations. **e**, Same as **b** but for the average of V1 intra-hemispheric correlations. *(**a–e**) See Table S1 for details about the statistics. See Figure S1 for the number of mice at each time point*.

In contrast, locomotion revealed a more pronounced reorganisation. Connectivity reductions were largely confined to the visual cortex (Fig. 8a), accompanied by a rapid and persistent decrease in network similarity across the entire recording period (Fig. 8b). Homotopic correlations were selectively reduced within the visual cortex (Fig. 8c), consistent with the observed suppression of visual activity during movement. V1 intra-hemispheric correlations decreased with posterior regions (Fig. 8d) and declined overall from day 1 to week 10 (Fig. 8e).

**Fig. 8.**
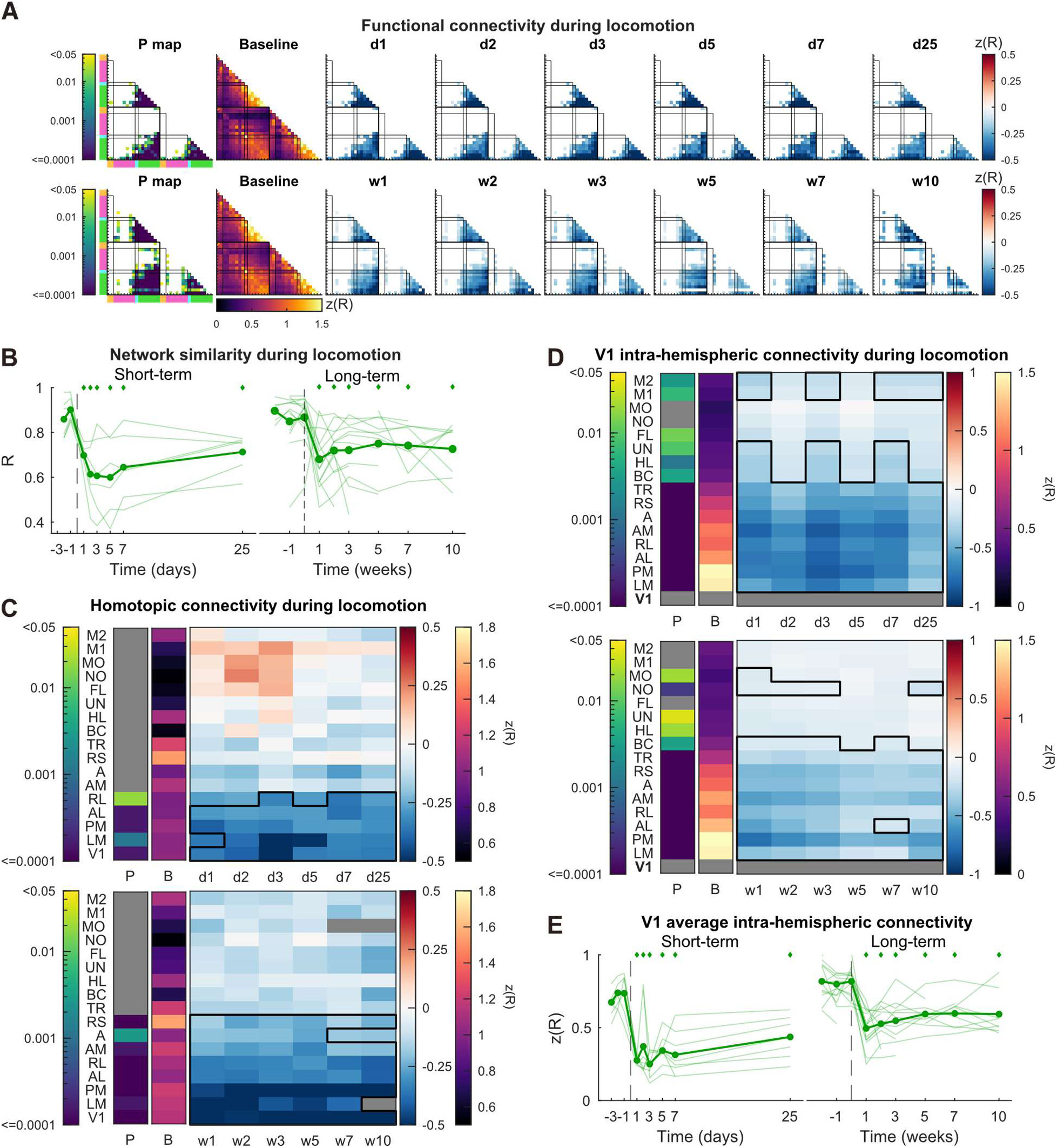
Reorganisation of the locomotion cortical network. a, Functional connectivity matrices calculated on the calcium signal (ΔF/F_0_) during locomotion using the Pearson correlation coefficient corrected with the Fisher z-transformation (z(R)). In the P matrices, only p values < 0.05 (linear mixed-effects model ANOVA, FDR-adjusted across pairs) are shown with a log10-scaled colour bar. The baseline matrices are the average correlation between mice and baseline sessions. The matrices from time points after BE show the difference from baseline, with only connections with a significant time effect shown. **b,** Network similarity. Thin lines represent individual mouse data. Diamonds mark significant time points relative to baseline (*p* < 0.05, post hoc FDR-adjusted). c, Homotopic correlations across cortical regions and time. *P* matrices (P) indicate regions with significant effects of time in a linear mixed-effects ANOVA, FDR-adjusted across regions (*p* < 0.05). The colour bar indicates log10(*p*). Baseline matrices (B) show the correlations averaged across mice and sessions. Post-BE matrices show the difference from baseline. The black outline indicates time points and regions that are significantly different from baseline (*p* < 0.05, post hoc FDR-adjusted). **d**, Same as **b** but for the V1 intra-hemispheric correlations. **e**, same as **b** but for the average of V1 intra-hemispheric correlations. *(**a–e**) See Table S1 for details about the statistics. See Figure S1 for the number of mice at each time point*.

Together, these results demonstrate that BE induces a rapid, long-lasting, and behaviour-dependent reorganisation of cortical network structure, with stronger effects during locomotion than quiescence.

## Discussion

This longitudinal exploratory study revealed month-long plasticity of cortical states following adult vision loss. We identified two sequential and overlapping windows of plasticity defined by behaviours, with spatially distinct effects on the cortex. The first one, from day 1 to week 3-5, was characterised by a reduction in visual cortex activity during locomotion and a suppressive effect of movements. This effect was observed in the visual and retrosplenial cortex. The second one, from week 1 to week 7-10, was characterised by an increase in slow-wave activity during quiescence and a rapid increase in activity at the transition from locomotion to quiescence. It was localised in V1 and LM and suggests increased excitability. These behaviour-dependent changes in activity induced by BE were accompanied by a rapid and long-lasting cortical network reorganisation, which was more extensive during locomotion than during quiescence (Fig. 9).

**Fig. 9.**
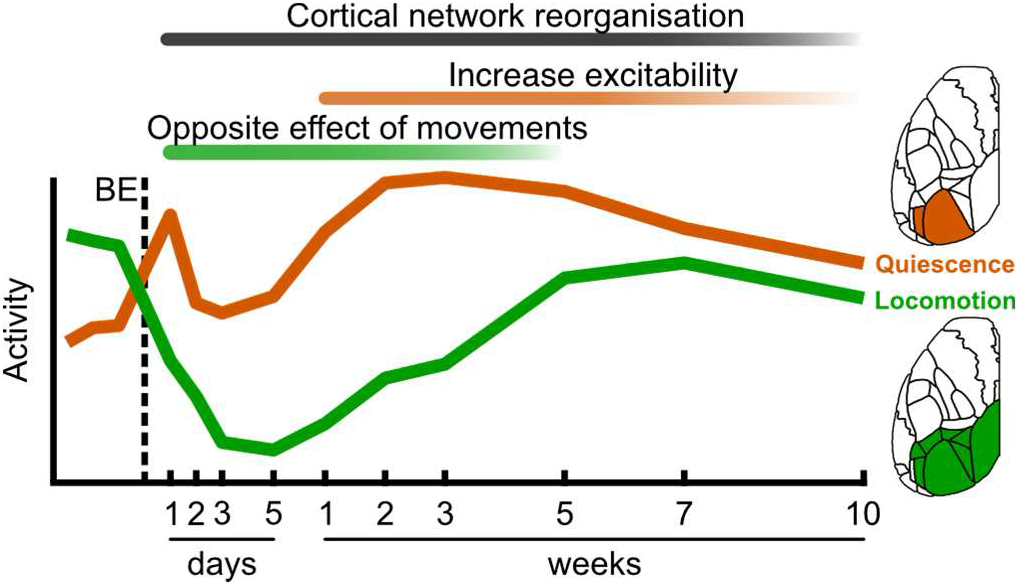
**Summary of main findings.**

### Plasticity and cortical states

Our results suggest long-term plasticity in the circuit regulating cortical states. In normal adult mice, locomotion desynchronizes the visual cortex spontaneous activity (Polack et al., 2013; Reimer et al., 2014; Vinck et al., 2015) and increases the gain of visual responses while maintaining their selectivity (Niell & Stryker, 2010). In the mesoscale calcium signal, this effect is evident as a sustained increase in fluorescence (West et al., 2022), as seen during our baseline recordings. This cortical state change is thought to be mediated by a disinhibitory circuit (Fu et al., 2014). When mice run, cholinergic neurons in the basal forebrain are activated, which in turn activate VIP inhibitory neurons (Alitto & Dan, 2013; A. M. Lee et al., 2014). VIP interneurons are known to suppress mainly SST interneurons (Muñoz et al., 2017; Pfeffer et al., 2013), leading to a disinhibition of excitatory neurons. This circuit has already been suggested to have a role in adult cortical plasticity (Stryker, 2014). In adult mice, after early-onset, long-term monocular deprivation, visual stimuli presented through the deprived eye during locomotion recovered their cortical responses, whereas other stimuli did not (Kaneko & Stryker, 2014).

Activating VIP cells or silencing SST cells recapitulated the effect of locomotion on visual cortex plasticity (Fu et al., 2015). Therefore, it suggests that locomotion, through the VIP-SST disinhibitory circuit, enhances adult cortical plasticity. Our study adds to this by showing that cortical states undergo long-term plasticity in adults.

### Early window of opposite effect of movements on cortical activity

The observed plasticity of cortical states is initiated from the very first day of recording, with movements suppressing rather than increasing activity of the visual and retrosplenial cortex. This rapid reversal in the effect of locomotion can be caused by a reorganisation of the cortical circuitry. It could be mediated by homeostatic plasticity mechanisms, which are rapidly engaged to maintain excitatory activity around set points (W. Wen & Turrigiano, 2024). An important homeostatic mechanism is a reduction in inhibition that helps restore excitatory activity levels.

The activity of inhibitory neurons decreases 12h after monocular enucleation but does not recover in the subsequent days, in contrast to the excitatory neurons. A structural disengagement of inhibitory neurons in the network mediates this sustained reduction in inhibitory activity. At day 2, the density of excitatory dendritic spines on inhibitory neurons decreases in V1, suggesting that they are less excited by excitatory neurons (J. L. Chen et al., 2011; Keck et al., 2011). Additionally, there are fewer inhibitory synaptic boutons on excitatory neurons, directly reducing the potential inhibition of their activity (Keck et al., 2011, 2013). Therefore, there is a profound reorganisation of the neuronal network structure that favours the stabilisation of excitatory activity after visual deprivation. This is likely to change the cortical circuit regulating cortical states, as locomotion’s effect on pyramidal neuron activity depends on a specific disinhibitory circuit. The modulation is maintained from day 1 to week 5, as indicated by differences between quiescence and locomotion, suggesting that, despite the circuit reorganisation, a pathway remains linking movements to pyramidal neuron activity, but in the opposite manner to what is typically observed. Interestingly, at weeks 7 and 10, there was no difference between continuous epochs of quiescence and locomotion activity, but an effect of movements was still present at the transition. This dissociation between the acute and sustained effects of the cortical state change might reflect distinct circuit components.

Cortical states are also controlled by the thalamus (Poulet & Crochet, 2019). Pharmacological inactivation of the thalamus has been shown to alter cortical states related to movement in the somatosensory (Poulet et al., 2012) and visual cortices (Nestvogel & McCormick, 2022). In V1, the depolarisation of the membrane potential of excitatory neurons normally observed following whisking and locomotion onset is reduced after thalamic inactivation. At whisking onset, the initial rapid depolarisation remained, but the amplitude was reduced. However, the sustained component that follows was completely abolished (Nestvogel & McCormick, 2022). Even though the modalities differ, we observed a similar effect after BE in the calcium imaging signal (Fig. 4). This suggests that, in addition to plasticity in cortical circuitry, thalamocortical plasticity could contribute to the reversal of movements’ effects on cortical activity after BE. Structural and functional plasticity has been observed in thalamocortical neurons after BE in adults (Bhandari et al., 2022), but more evidence is needed to better understand the role of the thalamus in adult cortical plasticity.

### Delayed window of increased cortical excitability

We identified a window from week 1 to week 7-10 during which the visual cortex shows increased slow-wave activity only when mice were quiescent, suggesting a state of increased excitability.

Increased excitability or responsiveness of the visual cortex has been observed following different types of visual deprivation in adults. In protocols of ocular dominance plasticity, the open-eye response increased only when the monocular deprivation lasted at least 7 days (Sato and Stryker, 2008). This is similar to juveniles, but they rely on different plasticity mechanisms. In mice lacking synaptic scaling, an important homeostatic plasticity mechanism, the open-eye potentiation was absent in juveniles but conserved in adults. However, in mice lacking the autophosphorylatable αCaMKII, this effect was absent in adults (Ranson et al., 2012). The autophosphorylation of αCaMKII is an essential process for the proper expression of adult long-term potentiation (Kirkwood et al., 1997), suggesting that this synaptic plasticity mechanism might be involved in the delayed increase in cortical responsiveness. A similar effect has been observed following monocular retinal lesions. In cats, spontaneous activity in the lesion projection zone of the visual cortex increases relative to the surround area at 2 and 4 weeks following a lesion, then recovers after 12 weeks (Giannikopoulos & Eysel, 2006). This is accompanied by an increase in αCaMKII autophosphorylation (Van den bergh et al., 2003), suggesting similar plasticity mechanisms to those observed during ocular dominance plasticity. Despite different deprivation methods and species, the period of increased spontaneous activity matches well with our data. Altogether, it suggests that the delayed increase in cortical excitability is a fundamental mechanism of adult cortical plasticity across species, which seems mediated by Hebbian plasticity mechanisms. This hyperexcitable state has been hypothesised to facilitate plasticity (Eysel & Jancke, 2024), and our study shows that it is behaviourally gated and persists for several weeks.

Our data showed that the increase in spontaneous activity was specific to the slow band. Cortical slow oscillations are known to mostly occur during sleep and anaesthesia, but can emerge locally during wakefulness after prolonged waking (Vyazovskiy et al., 2011), around cortical lesions (Sarasso et al., 2020), and when the thalamus is silenced (Nestvogel & McCormick, 2022; Poulet et al., 2012). These oscillations are characterised by a sequence of high firing (ON-period) followed by a period of hyperpolarisation (OFF-period) (Massimini et al., 2024; Steriade et al., 1993). During sleep, cortical slow waves have been shown to promote long-term plasticity (Chauvette et al., 2012; Timofeev & Chauvette, 2017). Therefore, it would be possible that, following BE in adults, the visual cortex “falls asleep” during quiescence to help reorganise cortical circuitry. However, it remains to be tested whether the slow oscillations detected in the calcium signal share the same electrophysiological signatures as those found during slow-wave sleep.

Prior to the window of increased excitability during quiescence, we found a transient increase in activity at day 1 after BE. This peak of activity was not found using two-photon calcium imaging of excitatory neurons following complete bilateral retinal lesion (Keck et al., 2013) or monocular enucleation (Barnes et al., 2015). However, the correlation between excitatory neurons was transiently increased at day 1, whereas activity was not, following BE (Mesik et al., 2026). With mesoscale calcium imaging, we mostly measure the correlated activity of large groups of neurons, as the signal likely arises from the sum of fluorescence from cell bodies and the neuropil (Barson et al., 2020; Cardin et al., 2020). Therefore, the change in correlation between excitatory neurons at day 1 can explain the transient increase we observed. It might result from homeostatic plasticity mechanisms, specifically synaptic scaling in excitatory neurons that occurs after visual deprivation to restore excitatory activity levels (Hengen et al., 2013; Keck et al., 2013; G. Turrigiano, 2011; G. G. Turrigiano et al., 1998).

### Long-lasting reorganisation of the cortical network

The cortical networks reorganised rapidly and in a behaviour-dependent manner. This is likely due to the localised change in activity found in each behavioural state. But over time, the cortical network changes do not show the same trajectory as the activity. At week 10, the activity levels partially recovered, whereas network measures such as network similarity and V1 intra-hemispheric connectivity did not. It suggests that the cortical functional connectivity can evolve independently of activity levels. While activity levels recovery suggests a long-term stabilisation of the activity regime, the fact that the correlation with other regions did not recover suggests that the information contained in the activity has not recovered. Long-term changes in functional connectivity are also observed in late-blind individuals (Z. Wen et al., 2018), suggesting that the network structure may never fully recover. Monocular deprivation during the critical period also reorganises the cortical network in the first 3 days. Notably, there is an increase in V1 connectivity with the parietal cortex and a decrease in V1 homotopic connectivity (S. Chen et al., 2024). Interestingly, the changes we found seem more profound, but it remains to be tested whether this is due to differences in developmental stage or in the visual deprivation method.

### Cortical state plasticity as a transition to cross-modal plasticity?

Interestingly, the gradual recovery of activity during locomotion observed from week 1 onward (Fig. 3d) parallels the time course reported by Van Brussel et al., 2011, who found a progressive restoration of V1 activity—assessed via the immediate early gene *zif268*—between 1 and 7 weeks following monocular enucleation. This method likely reflects an average of the activity in the last hour before brain sampling (Kaczmarek & Chaudhuri, 1997), a timescale necessary for gene transduction and mRNA accumulation in the cytoplasm. Although the mouse behaviour was not specified prior to sacrifice, our data would suggest that the mice were active. When the mice were also deprived of whiskers, activity recovery in V1 did not occur, suggesting that cross-modal plasticity underlies this recovery. We do not have evidence of cross-modal recruitment of the visual cortex, as this would require sensory stimulation (Hashimoto et al., 2023) or additional sensory deprivation (Van Brussel et al., 2011). However, we can hypothesise that it occurred several weeks after BE. It has been proposed that increased excitability following sensory deprivation could cause pre-existing subthreshold cross-modal inputs to reach the spiking threshold in the deprived cortex (Kral & Sharma, 2023). Activity during locomotion started to recover at the same time the activity increased during quiescence, at day 7. This state of increased excitability might strengthen cross-modal inputs, restoring activity levels during locomotion and partially restoring movement-evoked activity from week 5 onward, a period where activity during quiescence declined toward baseline levels.

## Materials and Methods

### Animals

All procedures were approved by the Animal Care Committee of the Université de Montréal (Comité de Déontologie en Expérimentation Animale, CDEA) and conformed to the guidelines of the Canadian Council on Animal Care and Use. We used 26 transgenic Thy1-jRGECO1a mice (GP8.20Dkim/J) that expressed the calcium indicator in glutamatergic pyramidal neurons of layer V and II/III of the cortex (Dana et al., 2018). The mice were separated into two main protocols, one over days (short-term) and one over weeks (long-term) after BE. The short-term protocol included 8 mice (4F/4M) aged 114-126 days at the time of BE. The long-term protocol comprised 2 cohorts, with variations detailed below. Experiments on cohort 2 started after cohort 1. All mice from a given cohort were recorded during the same period and using the same protocol. Cohort 1 included 10 mice (6F/4M) aged 162–198 days at the time of BE. Cohort 2 included 8 mice (3F/5M) aged 82–83 days at the time of the BE. The mice were housed in an inverse light cycle room (12h-12h) in groups of 2 or more, when possible, with only 2 males starting the protocol alone in a cage. A wheel was placed in each cage for mice to get used to running on it. In each recording session, males were recorded first, followed by females. The order of cage passage was conserved throughout the protocols, but within each cage, the mice were selected at random.

### Implantation of the transcranial chronic imaging window

We induced anaesthesia using 3% isoflurane in pure O2 and then maintained it at 1.5%. Once anaesthetised, mice were head-fixed on a stereotaxic frame. The eyes were protected with an eye lubricant, and the body temperature was maintained at 37 °C using a rectal probe connected to a feedback-regulated heating pad under the mouse. We removed the fur on the scalp and cleaned the skin with 70% ethanol and 10% providone-iodine using cotton-tipped swabs. We then injected the anti-inflammatory agent Carprofen (0.01mL/g) subcutaneously and the local anaesthetic Lidocaine under the scalp (0.05mL). The scalp skin was removed using scissors to match the shape of the future implanted chronic window, and the skull was cleaned with sterile cotton-tipped swabs to remove all remaining tissues and fluids. The cut skin surrounding the future implant site was secured with a tissue adhesive (Vetbond, 3M). As part of the dorsal skull is now exposed, we implanted a window (cover glass, hand-cut to a diameter of ∼9mm) over the dorsal cortex and a head bar (3 x 3 x 30 mm, titanium) behind, both of which were glued with clear dental cement (C&B Metabond). Once dry, we injected saline subcutaneously and stopped the anaesthesia. The mice were allowed to recover for 5 days post-surgery, including 2 days of monitoring and administration of Carprofen (0.01 mL/g).

### Bilateral enucleation in adult mice

The procedure involves ketamine anaesthesia for cohort 1 and isoflurane for cohort 2.

For Cohort 1, the procedure was similar to Jeroen Aerts et al., 2014. We anaesthetised the mice with a combination of ketamine (75 mg/kg) and dexmedetomidine (0.5 mg/kg). The eyes were disinfected with cotton-tipped swabs soaked in providone-iodine, and the lids were lifted from the orbit by pressing on the canthus with forceps. To locally anaesthetise the eyes, we applied Alcaïne for a few minutes. Then, on each eye, we pressed on the optic nerve with forceps and made circular movements until the eyeball was removed. Small pieces of absorbable hemostatic sponge (Spongostan) were inserted in the orbit to absorb the remaining fluids. To keep the eyelid closed, we used a tissue adhesive (Vetbond, 3M) to glue it. Anaesthesia was reversed with atipamezole (0.5 mg/kg), and the mice recovered in a heated cage with an oxygen supply. They received Carprofen injection (0.01 mL/g) for the next 2 days, and their health was monitored.

For cohort 2, we induced anaesthesia with 3% isoflurane and then maintained it at 2.5%. Similar to the imaging window implantation, the mice were placed on the stereotaxic frame, and their body temperature was maintained at 37 °C. We injected Carporfen subcutaneously (0.01mL/g). We removed the fur around the eyes and disinfected the area using cotton-tipped swabs soaked in providone-iodine. To remove each eye, we used forceps to press on the optic nerve and rotated around the forceps axis until the eyeball detached. Haemorrhages could happen after this step. In that case, we pressed on the orbit with a sterilised cotton tip. Once the haemorrhage stopped, we inserted small pieces of hemostatic sponge (Spongostan) into the orbit and closed the eyelid with tissue adhesive (Vetbond, 3M). At the end of the surgery, the mice received a saline injection and were then recovered in a heated cage. They received a Carprofen injection (0.01 mL/g) for the next 2 days, and their health was monitored.

### Longitudinal protocols

After implanting the imaging window and the recovery period, we began a week of gradual habituation to the head fixation system. On day 1, the mice were manipulated for 5 minutes. On day 2, they were manipulated and allowed to explore the fixation system, which included the running wheel. From Days 3 to 5, the mice were head-fixed for 5 to 10 minutes, with manipulation and exploration preceding this period. Once habituated, the longitudinal protocols began with 3 days of baseline sessions for the short-term protocol, 4 weeks for cohort 1, and 3 weeks for cohort 2 of the long-term protocol. During baseline, we performed retinotopic mapping of the visual cortex (see section below). We also conducted one session under anaesthesia (cohort 2 only), which included tactile stimulation of the left hindlimb, forelimb, and whiskers to map somatosensory areas (not presented in this article, see section below). After completing the baseline sessions, all the mice underwent BE. In the short-term protocol, we recorded spontaneous cortical activity from day 1 after BE. We maintained the same order of passage between surgeries and imaging sessions, so that the mice were imaged at 24h±2h after BE. The following measurements were made at 2, 3, 5, 7, and 25 days after BE. In the long-term protocol, the awake spontaneous cortical activity was recorded at 1, 2, 3, 5, 7, and 10 weeks after BE. Additionally, with cohort 2, we performed sessions under anaesthesia at 1, 3, 5, and 10 weeks after BE (not presented in this article).

### Mesoscopic imaging

We used mesoscopic calcium imaging of the adult mouse dorsal cortex, expressing the red calcium indicator jRGECO1a in excitatory neurons. For spontaneous and tactile stimulation sessions, we used a dual-camera system from Labeo Technologies Inc. to capture the calcium and hemodynamic signals. For both cameras, images of 160 × 160 pixels were acquired at 60 Hz with a field of view of ∼10.5×10.5 mm. The system had three LEDs: lime (565/24 nm) for jRGECO1a excitation, and green (525 nm) and red (625 nm) for the hemodynamic measurements. Camera 1 captured the red fluorescence from the calcium indicator and the red reflectance from the red LED (filter 618/50), and camera 2 captured the green reflectance from the green LED (filter 514/44nm). The light from the cortex was directed to camera 2 using a dichroic mirror (Di552nm). Cameras 1 and 2 captured a first frame when the lime and green LEDs were on, while the red LED was off. In the next frame, the red LED was on alone and captured by camera 1 and so on. Therefore, each colour channel was sampled at 30Hz.

For the retinotopic mapping sessions, we used a single camera mesoscope from Labeo Technologies Inc. There was a lime (565/24 nm) LED for jRGECO1a excitation and a red emission filter (618/50 nm) to capture the red fluorescence. Images of 1024×512 pixels were acquired at 10Hz with a field of view of ∼10.5×5.25 mm.

### Behavioural monitoring

During the awake spontaneous sessions, the mice were free to run on a 16.5 cm diameter wheel tilted upward by 15° (de Vries et al., 2020). The wheel was made from 3D printed parts and covered with a non-abrasive grip tape. We measured the mice’s behaviour using a camera under infrared illumination. Images of 1024×600 binned 2x were acquired at 30Hz.

### Retinotopic mapping

We mapped the visual areas using the retinotopic mapping method previously described (Kalatsky & Stryker, 2003; Zhuang et al., 2017). Briefly, we positioned a monitor (cohort 1: 24”, cohort 2: 27”) at 13.5 cm in front of the mouse’s eye, tilted 20° from the vertical axis, and rotated 30° from the mouse’s dorsoventral axis. A 20° bar containing a chequerboard swept in the four cardinal directions, with 10 repetitions for each direction (similar parameters as Zhuang et al., 2017). Each eye was mapped 2-3 times on different days during baseline.

### Awake sessions

The cortex’s spontaneous activity was measured while the mice were free to run on a wheel. Before the experiment, mice had 5 min to explore the experimental rig freely. Then, they were head-fixed on the running wheel. The implant was cleaned using 70% ethanol applied with cotton-tipped swabs, and if necessary, some fur could be trimmed to ensure a clear optical path to the cortex. To reduce the amount of light from the LEDs that could reach the eyes, a custom-made 3D-printed light cover was positioned around the implant. After placing the mouse under the mesoscope, spontaneous sessions were conducted for 20 min in the short-term and cohort 2 of the long-term protocol. For cohort 1, it consisted of two 10 min sessions, conducted consecutively over two days.

### Processing mesoscale cortical imaging

The views from the dual-camera imaging system were slightly offset. Camera 1 captured the red channels, and camera 2 captured the green channel. To align camera 2 on camera 1, we imaged a 1951 USAF resolution target. We used a local feature detection (detectKAZEFeatures in MATLAB) and an extraction and matching algorithm to find matching points between images (will be referred to as KAZE-based registration). From those points, we computed a 2D geometric similarity transformation. This transformation was then applied to the green frames from every imaging session on the dual-camera system.

The Allen Mouse Brain Atlas (Oh et al., 2014) was first placed on a reference spontaneous session, selected from the baseline sessions for each mouse. For precise placement, we used functional maps. Visual areas from both hemispheres were identified using retinotopic mapping and the phase-based analysis method previously described (Kalatsky & Stryker, 2003; Zhuang et al., 2017). One or more retinotopic mapping sessions have been performed on each eye on different days, therefore requiring alignment. We used the KAZE-based registration followed by a 2D geometric similarity transformation. Once aligned, the visual maps were combined to form a single map. For mice that underwent anaesthetised sessions, we identified somatosensory areas of the whiskers, forelimb, and hindlimb. We trial-averaged the evoked tactile response at 0.5s after stimulation, normalised using the 1s pre-stimulation period as the baseline. The resulting maps were directly combined in a single somatosensory map, as alignment was not necessary because all tactile stimulations were conducted in the same imaging session. Then, the visual and somatosensory maps were aligned on the reference spontaneous session using the KAZE-based registration followed by the 2D geometric similarity transformation. We manually placed the atlas (ROImanager from the umIT toolbox, Souza et al., 2025) on a composite image so that visual and somatosensory areas, along with the cortical midline, matched as closely as possible.

After manually placing the atlas on one reference session for each mouse using functional and anatomical landmarks, we automatically applied it to all other sessions using the KAZE-based registration followed by the 2D geometric similarity transformation. The algorithm was mainly successful, but in rare cases of incorrect registration, we manually identified matching points between images and then applied the same 2D geometric similarity transformation. After this process, the atlas was positioned in every session; however, they were not aligned with each other.

Finally, we defined a global reference, which was a centred binary image of the atlas. To align every session to the global reference, we calculated the geometric similarity transformation using phase correlation (imregcorr in MATLAB) between binary images of the atlas. We then applied the transformation to every frame of every imaging session. After this step, all sessions and colour channels were aligned.

A cortical mask was drawn to select usable pixels over the cortex and remove artefacts from the surgery. In the long-term protocol, the mask was drawn for each mouse at every session to account for implant degradation that accumulated over time. In the short-term protocol, the mask was drawn once on the reference session (Fig. S8).

All imaging channels were parcellated using the mouse cortical atlas. For each region, pixels within a 0.25mm radius of the centroid were averaged. Only pixels inside the cortical mask and with a mean intensity of at least 3000 over 65535 were averaged at each frame. If more than 75% of pixels did not respect this criterion, the region was excluded. Therefore, at the end of this step, we had one calcium time series per cortical region.

Red calcium indicators are less sensitive to hemodynamic fluctuations than green ones, but they can still contribute slightly to the measured changes in fluorescence (Wang et al., 2024). To account for the possible contamination, the hemodynamic component of the calcium fluorescence was removed using linear regression of the green and red reflectance (Valley et al., 2020). Then all analyses were done on the corrected calcium signal.

During the spontaneous behaviour sessions, the mice occasionally moved their tails above the imaging window, contaminating the signal. To remove those events, we detected periods when the average calcium signal in the cortical mask exceeded 7 standard deviations. We extended these periods to 1s before and after. Then, for spectral analysis, these periods were filled with a predicted signal (fillgaps in MATLAB). Otherwise, it was filled with nans.

### Processing behaviour

To quantify the behaviour, we used Facemap to extract the motion energy (Syeda et al., 2024). We drew an ROI that included the forelimbs, hindlimbs, and the rest of the body below the face (Fig. 1d). The motion energy was calculated as the sum of the absolute differences between consecutive frames within the ROI. Because the camera view was kept consistent across the protocols, the ROI was placed once and used every subsequent session.

An initial rough estimate of locomotion periods was established by using the moving standard deviation (2s window) and smoothed motion energy (moving average with a 0.33s window) that exceeded a threshold. To precisely identify the start and stop transition times, we applied a change point detection algorithm, which detected changes in root-mean-square level (findchangepts in MATLAB) around each estimated transition. Sustained locomotion was defined as periods between locomotion onset and offset of at least 10s and separated from transitions by 3s. For state transition analysis, transitions were further selected to have no locomotion within the 3s preceding the start of locomotion and had continuous locomotion during the 5s following, and vice versa for locomotion offset (Vinck et al., 2015).

To identify quiescence periods, a process similar to that used for locomotion has been employed. Sustained quiescence was defined as periods between the onset and offset of at least 10s and separated from transitions by 3s.

During quiescence, we observed that the mice made small, brief movements that elicited a cortical response. To identify these micro-movements, we thresholded the smoothed motion energy (moving average with a 0.33s window) during quiescence. A change point detection algorithm was used to find the onset time. Only micro-movements with at least 5s without any movements before were kept.

### Data inclusion criteria

Considering the mice’s spontaneous behaviour in longitudinal protocols was challenging, as they did not necessarily express all measured behaviour at each time point (Fig. S1, S6). For example, a mouse could have run during the whole 20 min session, resulting in no data at that time point for quiescence and state transitions. Additionally, for a given mouse in the long-term protocol, we might have lost the signal of regions near the edge of the dorsal cortex because of implant degradation (Fig. S9). Therefore, for each region, behaviour and mouse, we included the data only if there was at least one baseline session and at least one session after BE. Moreover, each time point required data from at least three mice to be considered.

### Analysis of activity in sustained states

To quantify the activity levels for the correlation analysis (Fig. 2), we calculated the maximum power of the signal. For each region, the raw calcium fluctuations were normalised by calculating the ΔF/F_0_, which is the raw fluorescence minus the mean fluorescence, divided by the mean fluorescence. The continuous wavelet transform (CWT) of the ΔF/F_0_ was calculated using a symmetrical Morse wavelet with a time bandwidth of 1s. A Morse wavelet with these parameters is similar to a Morlet wavelet, but it was somewhat faster to compute. We extracted the power as the squared absolute values of the CWT. The output is a frequency-by-time matrix of power values. As a measure of ongoing activity, we calculated the maximum power values in the 0.1-4Hz band at every time point.

For the analysis of activity segmented by behavioural states (Fig. 3, S2), we calculated the power spectral density (pwelch in MATLAB) of ΔF/F_0_ for each behavioural-state block, using a 10s window (the minimum duration of a behavioural state) with 50% overlap. The result was a power spectrum per block. For each behavioural state, the power spectra were averaged. Activity levels were quantified as the maximum power values in the 0.2-4Hz band. Figures and statistics were generated using log10 power values.

### Analysis of activity at state transitions

For each state transition, we calculated the ΔF/F_0,_ using the average calcium signal from 5s before the transition as the baseline F_0_. All transition responses from the same type were averaged in a session. To compare the responses over time after BE, we calculated the average ΔF/F_0_ in the first second after the transition. To more precisely examine when the activity was affected relative to the transition time in V1, we showed the transition response over time from the transition and over time in the protocol. The transition response was binned 3x and was shown as the difference from the protocol baseline. To quantify the mouse movements from which state transitions were detected, we averaged the time-locked motion energy in the same window.

### Analysis of functional connectivity

The raw calcium signal was converted to ΔF/F_0_ and bandpass-filtered to 0.1-4Hz. The Pearson correlation coefficient was calculated between all pairs of regions during the whole session or during quiescence and locomotion periods. The Fisher z-transformation was applied to the correlation coefficients. From the correlation matrices, we calculated V1 intra-hemispheric connectivity as the average of connectivity within each hemisphere and homotopic connectivity for each region. We calculated network similarity as the correlation between each time-point matrix and the first baseline.

### Post-analysis processing

In cohort 1, two 10 min sessions were done per week. All analyses were done on each 10 min session. Then the data were weighted averaged to end up with one data per time point. The weights were the number of behavioural epochs.

The first baseline from cohort 1 (w-3) was removed as it was not present in cohort 2.

All analyses were done on every available cortical region. To simplify the results presentation, we averaged the data between hemispheres.

### Statistical analysis

For statistical tests, baseline sessions were averaged to create a single baseline time point. To account for missing data points from each mouse across the longitudinal protocols, we performed a linear mixed-effects model (lme from the nlme package in R) with time as a fixed effect and mice as a random effect. For the frequency power spectrum over time (Fig. 4b, S2c), frequencies were added as a second factor to the model. For the activity following state transition (Fig. 5b,d; 6b), the time from transition was added as a second factor to the model. For comparison between behavioural states, the state was added as a second factor to the model. To test whether there was an effect of time of BE, a two-sided ANOVA was performed on the model. For region data or connectivity matrices, the test was conducted independently for each region or pair of regions.

The p-values for the time effect were adjusted using the False Discovery Rate (FDR) method to account for multiple comparisons across regions. We concluded that there was a significant effect of time of BE when the adjusted p-value was < 0.05. As a post hoc test, we performed pairwise comparisons of the estimated marginal means between time points, with p-values FDR-adjusted. For a given measure and behaviour, the p-values of the post hoc from all significant regions were collected, and FDR adjusted a second time. Results from the statistical tests are presented in Table S1. The number of mice included in each region, behaviour, and time point is shown in Figures S1 and S6.

## Acknowledgments

We thank the following funding sources : Fonds de recherche du Québec – Santé https://doi.org/10.69777/312876 (ID), Fonds de recherche du Québec – Santé https://doi.org/10.69777/269359 (MPV), Natural Sciences and Engineering Research Council of Canada (MPV), Vision Science Research Network (ID, MPV), Centre interdisciplinaire de recherche sur le cerveau et l’apprentissage (ID), L’École d’optométrie de l’Université de Montréal (ID), Études supérieures et postdoctorales de l’Université de Montréal (ID). We thank Guillaume Poulain for performing the surgeries and caring for the mice, and the lab of Ravi Rungta for providing the transgenic mice. We thank Justine Zehr and Miguel Chagnon from the Department of Statistics at the Université de Montréal for their help in developing and validating our statistical approaches. We thank Elvire Vaucher, Greg Silasi, Sergio Crespo-Garcia, and Ravi Rungta for their thoughtful review of this manuscript and their constructive feedback. We used Grammarly to improve grammar and language quality throughout the manuscript.

## Author contributions

Conceptualisation: ID, MPV Methodology: ID, MPV Investigation: ID Visualisation: ID Supervision: MP, MPV Writing—original draft: ID

Writing—review & editing: ID, MP, MPV

## Code accessibility

The processed and analysed data, along with the code, are available in figshare with the identifier doi:10.6084/m9.figshare.30473159.

## Supplemental Material

Figures S1 to S8

Table S1. Statistics (separate file)

The table contains the results from the linear mixed-effects model ANOVA and post hoc tests for each figure.

**Figure S1.**
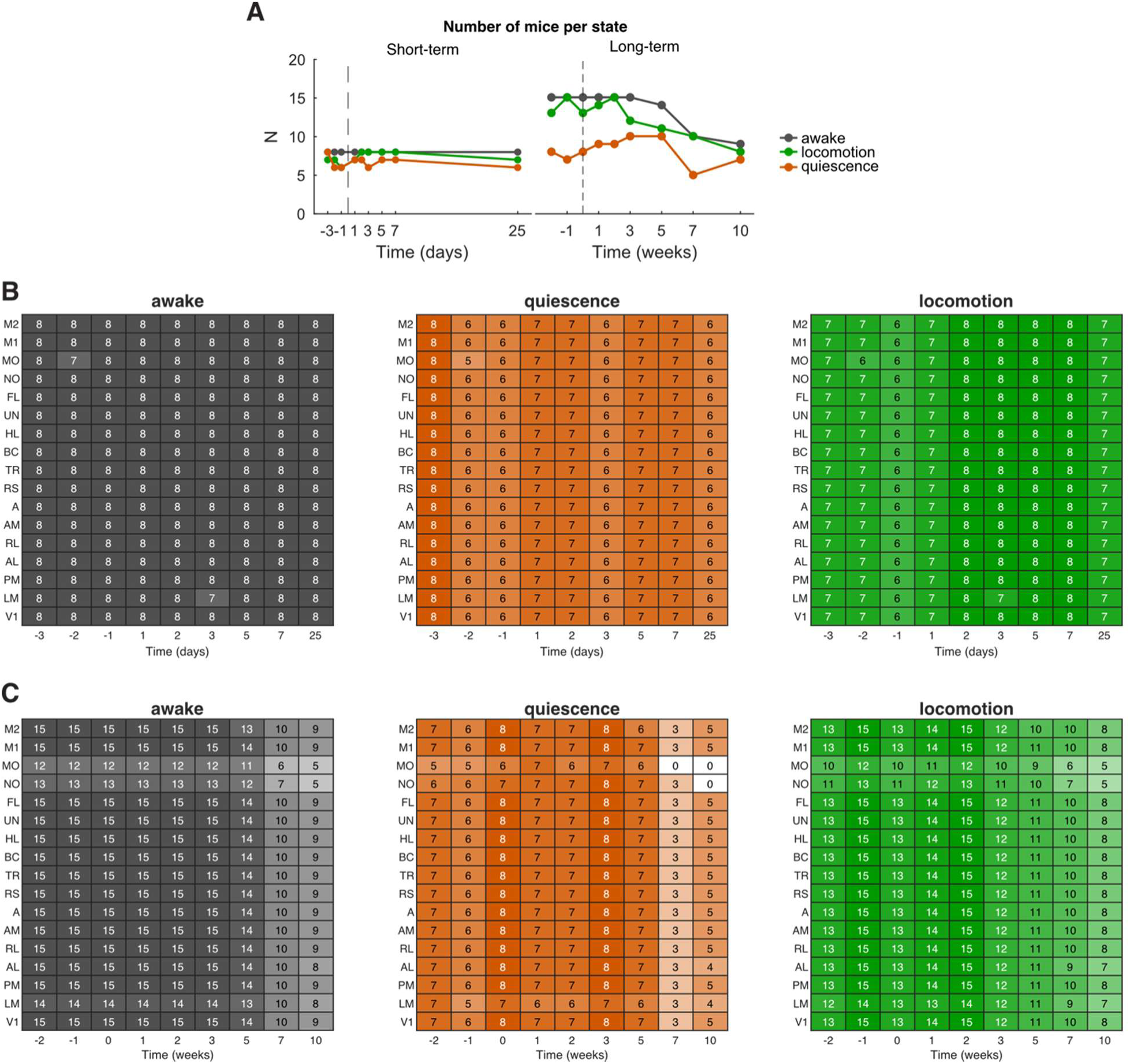
Number of mice in each state. **a** Number of mice in each state in both longitudinal protocols. As every mouse had at least one awake session per time point, the “awake” line represents the maximum number of mice at each time point. For locomotion and quiescence, the number of mice was equal to or inferior to the maximum (awake). Indeed, some mice had no quiescence and/or locomotion that entered the classification criteria. For a given mouse, this could also vary over time. **b** Number of mice in each state for each cortical region in the short-term protocol. Since regions were excluded based on the pixel intensity and implant quality (See Methods, Supplementary Fig. 5), the N for each region and behaviour is not consistent over time. Here, for each behavioural state, the heatmap shows the number of mice per region and time point. **c** Same as **b**, but for the long-term protocol.

**Figure S2.**
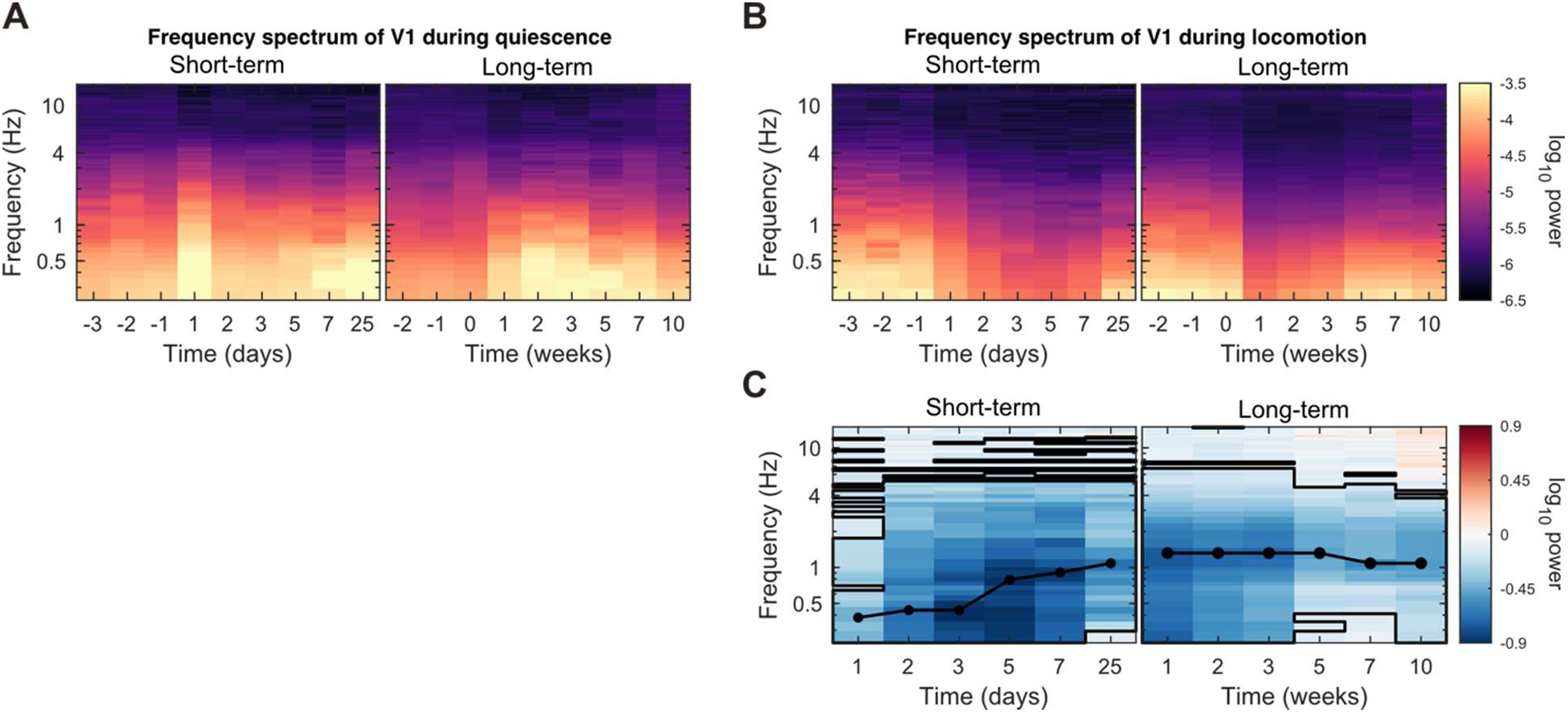
V1 power spectrums over time (related to. **Fig. 3). a** Power spectrum over time of the V1 activity during quiescence, shown in log10 power. **b** Same as **a** but for locomotion. **c** Frequency power spectrum of V1 activity during locomotion, shown as the difference with baseline. The black outline indicates time points and frequencies that are significantly different from baseline (p < 0.05, post hoc FDR-adjusted). *See Table S1 for details about the statistics. See Figure S1 for the number of mice included at each time point*.

**Figure S3.**
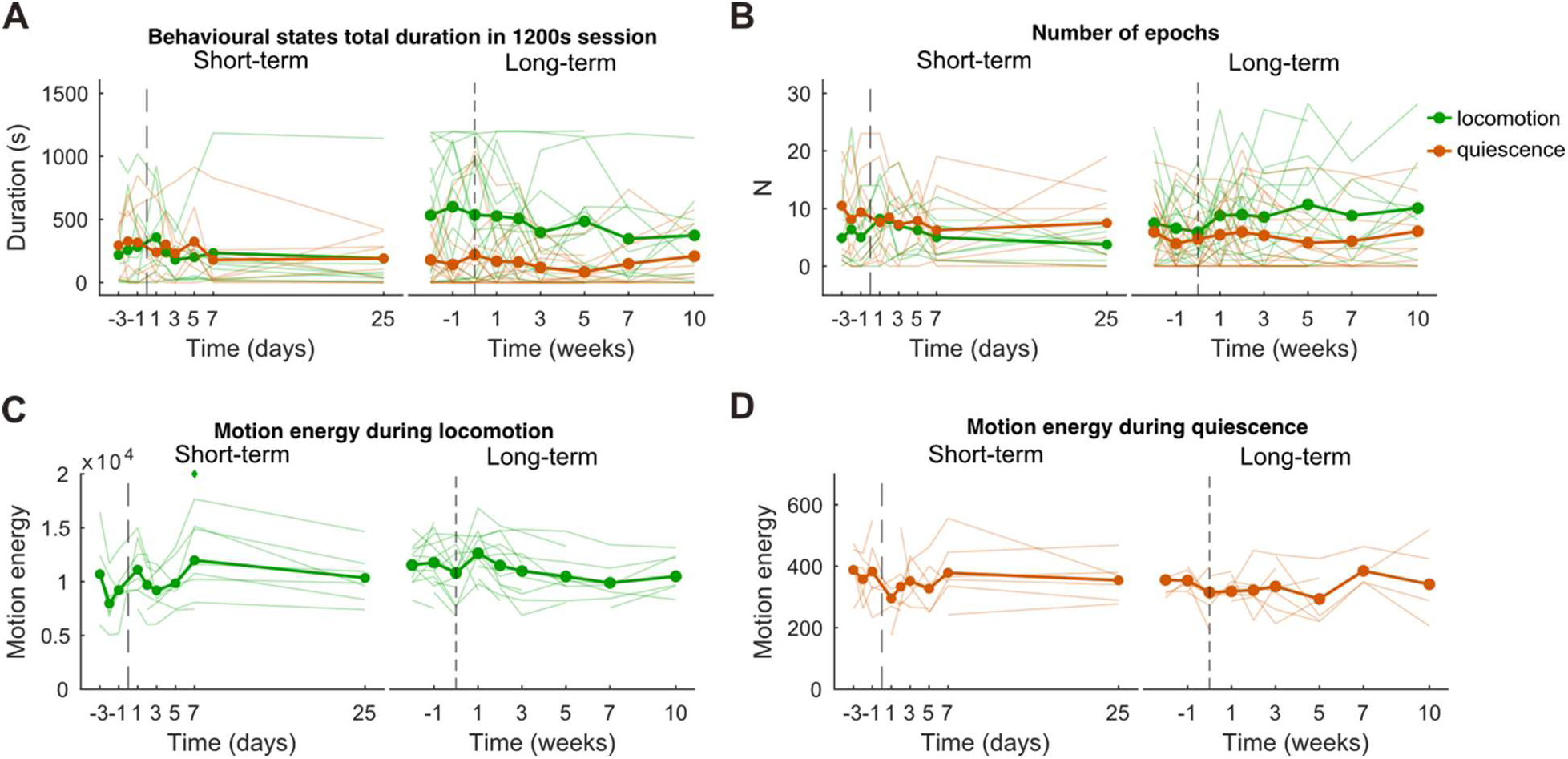
Behaviour over time. **a** Total behavioural state duration in a 20-minute session. **b** Number of behavioural state epochs. **c** Motion energy during locomotion. **d** Motion energy during quiescence. (**a-d**) Thin lines represent individual mouse data. Diamonds mark significant time points relative to baseline (*p* < 0.05, post hoc FDR-adjusted). *See Table S1 for details about the statistics. See Figure S1 for the number of mice included at each time point*.

**Figure S4.**
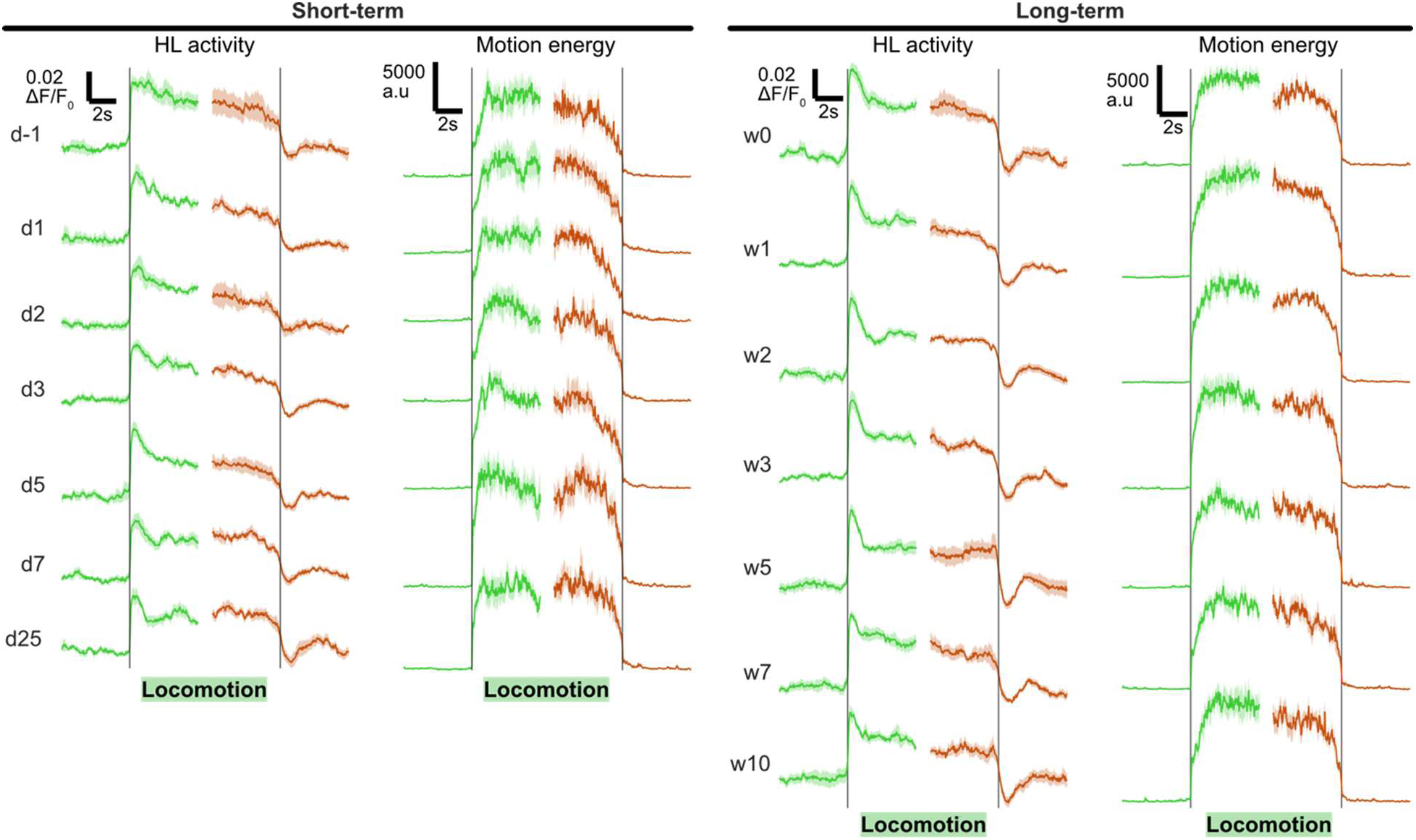
HL activity at the onset and offset of locomotion (related to. **Fig. 4). a** Mouse averaged ΔF/F_0_ response in HL and motion energy time locked on the locomotion onset and offset at one baseline and every time point after BE. The shaded areas represent the SEM.

**Figure S5.**
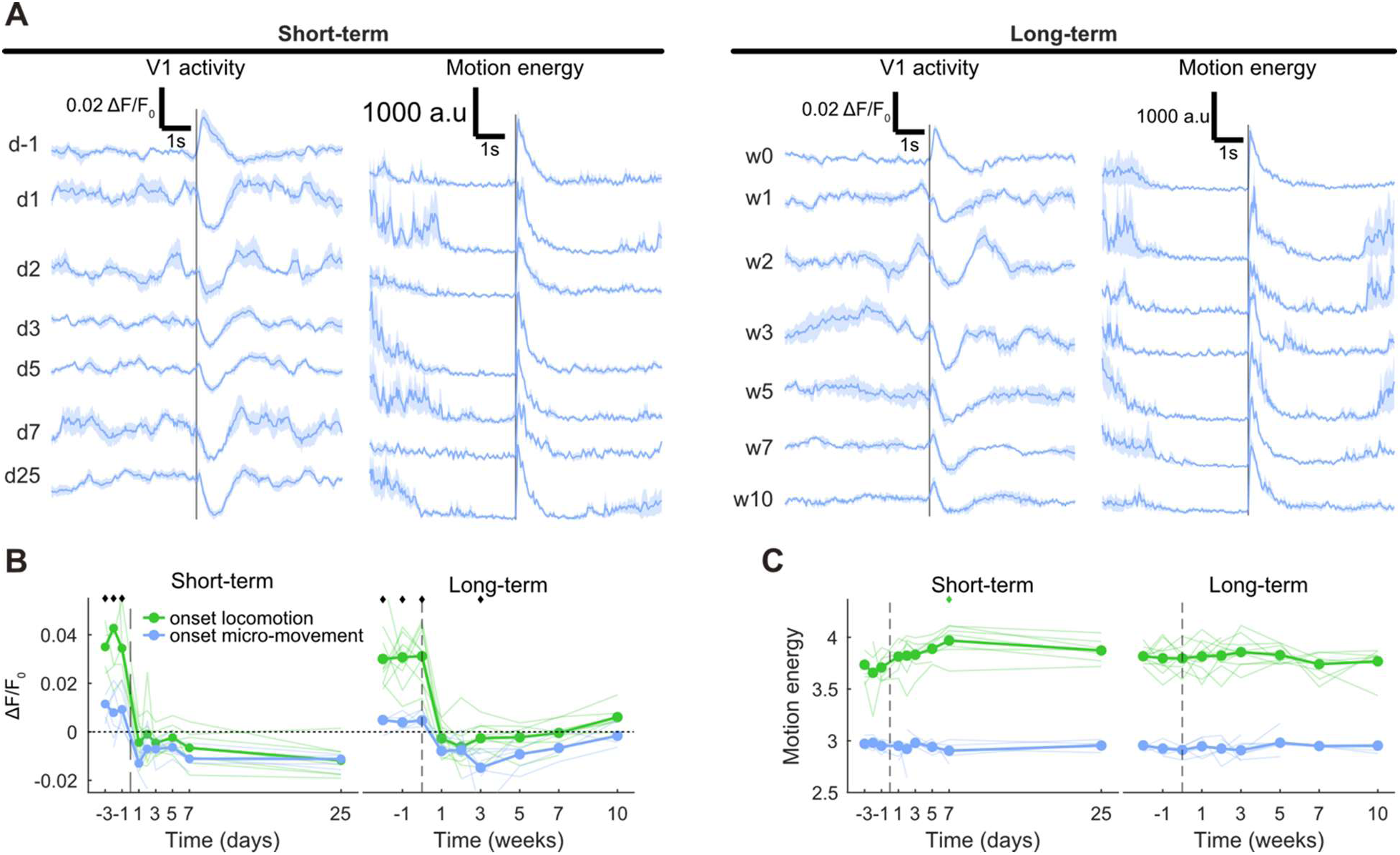
Related to. **Fig. 5. a** Mouse averaged ΔF/F_0_ response in HL and motion energy time locked on micro-movements at one baseline and every time point after BE. The shaded areas represent the SEM. **b** Average ΔF/F_0_ within a 1s window following micro-movements, normalised to the pre-transition period. Diamonds mark significant time points with differences between states (*p* < 0.05, post hoc FDR-adjusted). **c**, Average motion energy within a 1s window following micro-movements. Diamonds mark significant time points relative to baseline (*p* < 0.05, post hoc FDR-adjusted). (**b,c**) Thin lines represent individual mouse data. *See Table S1 for details about the statistics. See Figure S6 for the number of mice included at each time point*.

**Figure S6.**
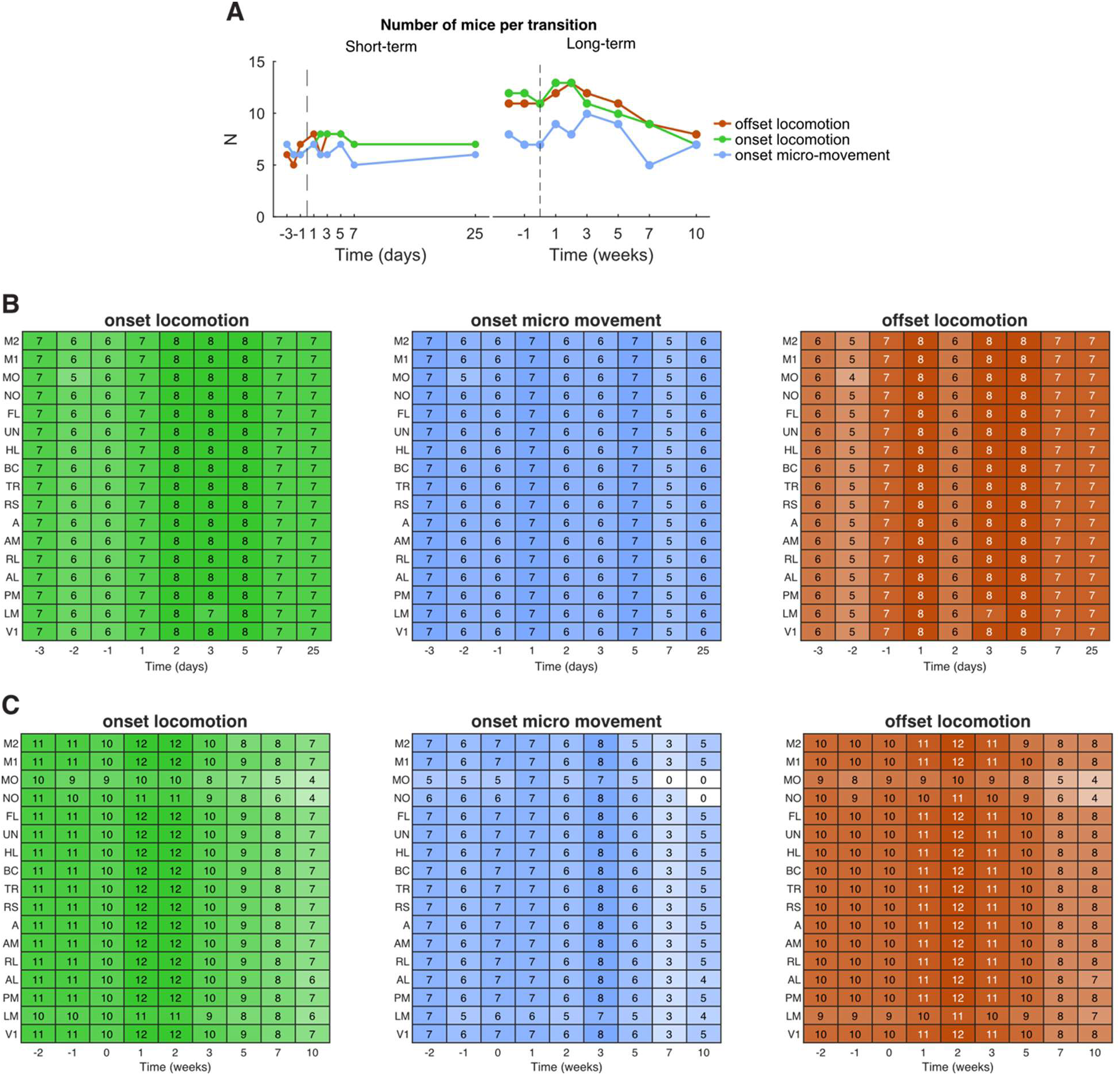
Number of mice in each state transition. **a** Number of mice per transition in both longitudinal protocols. The number of mice was equal to or less than awake in Supplementary Figure 1. **b** Number of mice in each transition for each cortical region in the short-term protocol. Since regions were excluded based on the pixel intensity and implant quality (See Methods, Supplementary Fig. 5), the N for each region and behaviour was not consistent over time. Here, for each behavioural state, the heatmap shows the number of mice per region and time point. **c** Same as **b**, but for the long-term protocol.

**Fig. S7.**
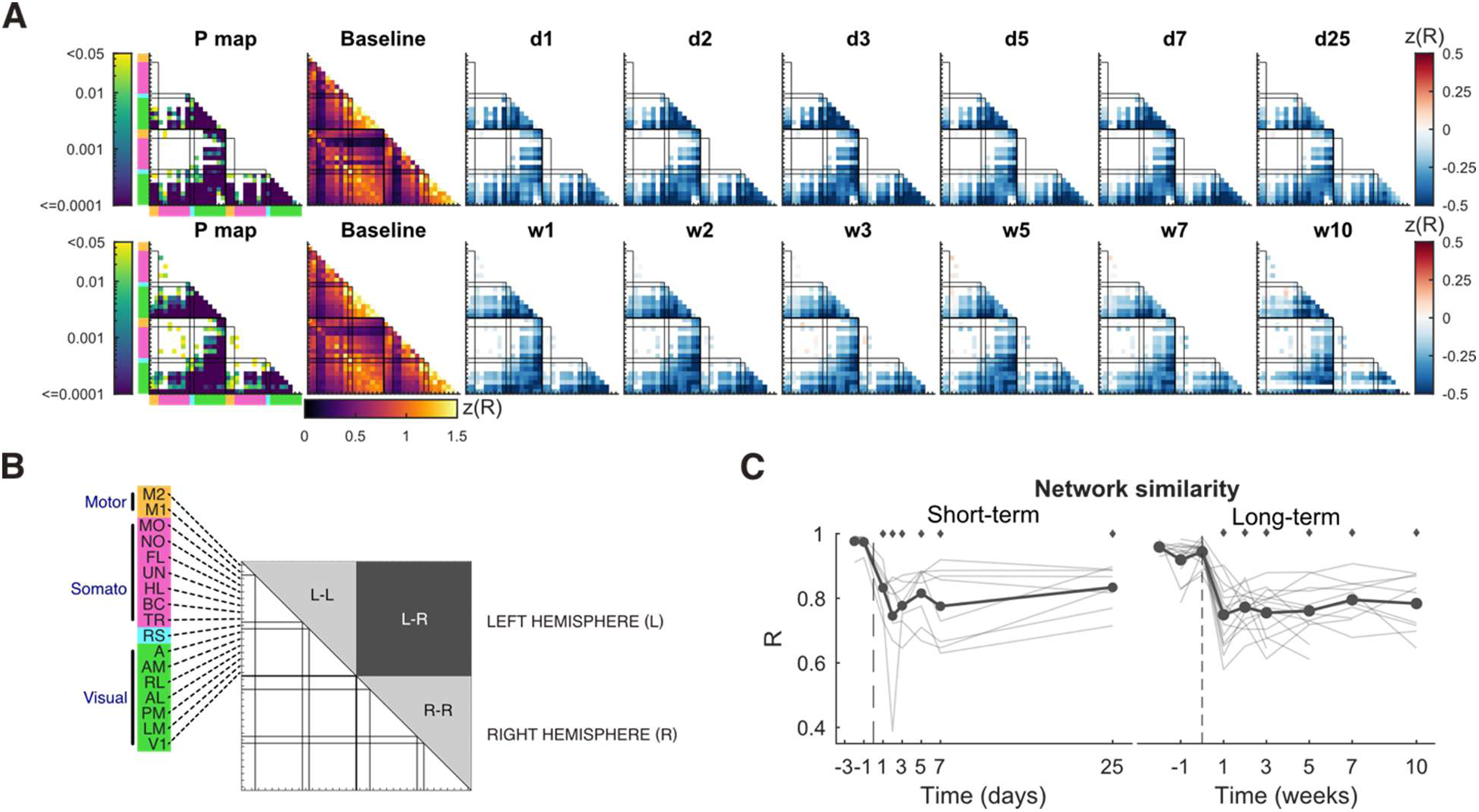
Cortical network reorganisation during spontaneous behaviour. **a** Functional connectivity matrices calculated on the calcium signal (ΔF/F_0_) during spontaneous behaviour using the Pearson correlation coefficient corrected with the Fisher z-transformation (z(R)). In the P matrices, only p values < 0.05 (linear mixed-effects model ANOVA, FDR-adjusted across pairs) are shown with a log10-scaled colour bar. The baseline matrices are the average correlation between mice and baseline sessions. The matrices from time points after BE show the difference from baseline, with only connections with a significant time effect shown. **b** Diagram of a connectivity matrix. In a hemisphere, regions are arranged in order according to their anteroposterior position. **c** Network similarity. Thin lines represent individual mouse data. Diamonds mark significant time points relative to baseline (*p* < 0.05, post hoc FDR-adjusted). *(**a–c**) See Table S1 for details about the statistics. See Figure S1 for the number of mice at each time point*.

**Figure S8.**
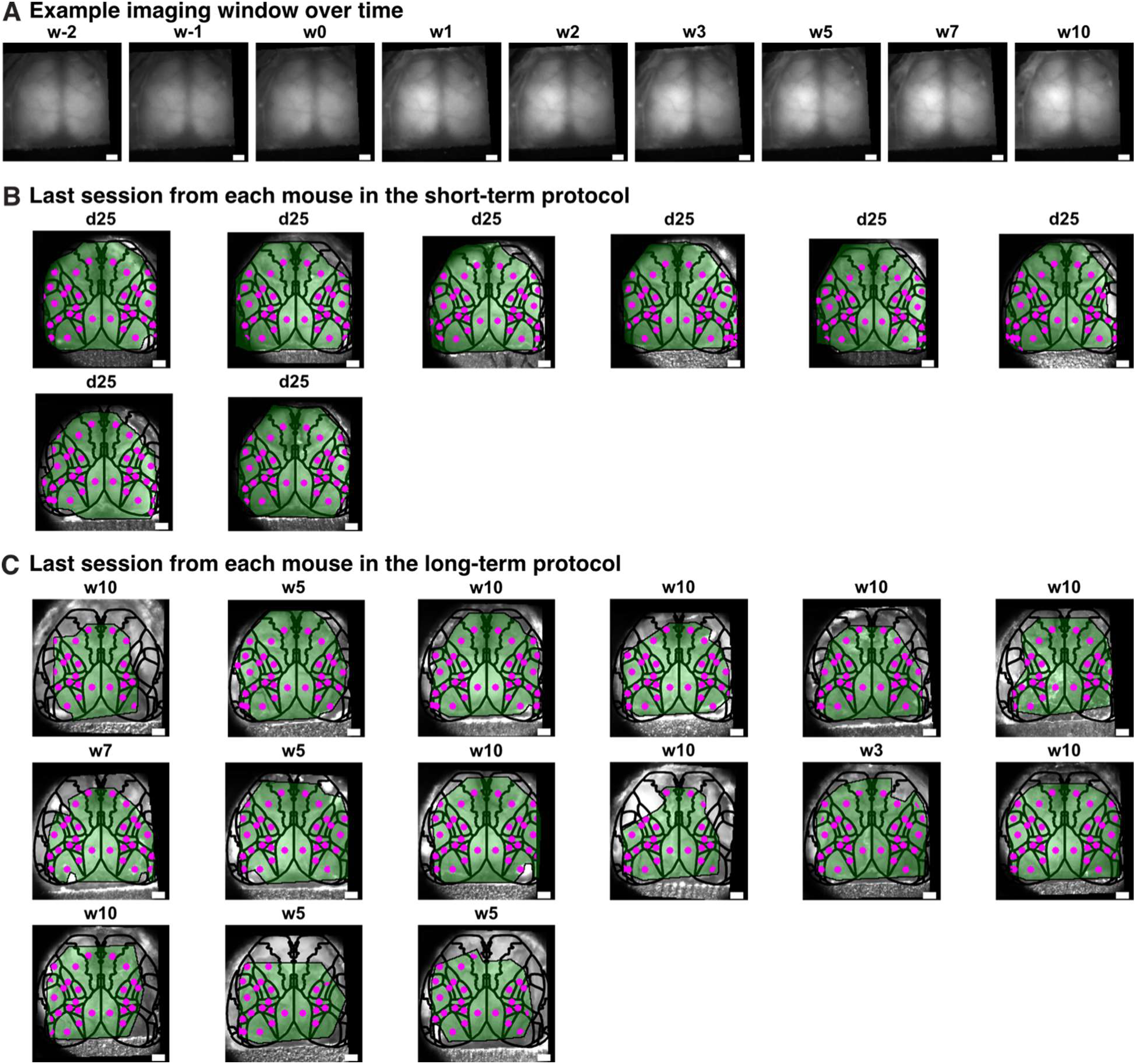
Chronic imaging window. **a** Imaging window viewed from the calcium channel over time from an example mouse. **b** Images from the green channel (green reflectance) showing the dorsal cortex seen through the chronic imaging window at the last session of each mouse. The atlas and the cortical mask are overlaid on it. In the short-term protocol, the cortical mask has been drawn once per mouse, on a reference session during baseline. **c** Same as **b**, but for the long-term protocol. The cortical mask has been drawn for each session of each mouse due to implant degradation over time. (**a-c**) Above each image, the corresponding time point is indicated. Scale bar is 1mm.

